# Testing for correlation between traits under directional evolution

**DOI:** 10.1101/566349

**Authors:** Manuela Royer-Carenzi, Gilles Didier

**Affiliations:** Aix Marseille Univ, CNRS, Centrale Marseille, I2M, Marseille, France; IMAG, Univ Montpellier, CNRS, Montpellier, France

**Keywords:** directional evolution, correlation tests, Brownian motion, independent contrasts

## Abstract

Being confounding factors, directional trends are likely to make two quantitative traits appear as spuriously correlated. By determining the probability distributions of independent contrasts when traits evolve following Brownian motions with linear trends, we show that the standard independent contrasts can not be used to test for correlation in this situation. We propose a multiple regression approach which corrects the bias caused by directional evolution.

We show that our approach is equivalent to performing a Phylogenetic Generalized Least Squares (PGLS) analysis with tip times as covariables by providing a new and more general proof of the equivalence between PGLS and independent contrasts methods.

Our approach is assessed and compared with three previous correlation tests on data simulated in various situations and overall outperforms all the other methods. The approach is next illustrated on a real dataset to test for correlation between hominin cranial capacity and body mass.

## 1 Introduction

Testing for correlation between two traits is a natural question which has been widely studied, notably in a comparative biology context (Groussin and Gouy 2011; Marchini et al. 2014; Grabowski et al. 2015; Will et al. 2017; Zhao et al. 2017). Correlation tests may concern any kind of traits: phenotypic, genetic or other. For instance, Seligmann (2018, 2019) observed a correlation between the syntheny level of poxviruses with amoeban mitogenome and their genome size. Assessing the correlation between two traits measured on several species cannot be performed by directly computing the Pearson correlation coefficient on the traits values since these values are not independent but related through the evolutionary relationships of the species involved (Diaz-Uriarte and Garland 1996). This point raises questions about how to interpret a correlation between two traits in a phylogenetic context. Actually, since all the observed taxa are assumed to have evolved from a single ancestral taxa, stating that two traits shared by the taxa are correlated can have only one significance, which is that the respective evolutions of these traits are correlated one with the other (Harvey and Pagel 1991). Assumptions about traits evolution are thus essential in order to disentangle the dependency structure of their extant values, and eventually to be able to study their correlation by correcting biases due to their evolutionary relationships (Martins and Garland 1991; Garland and Adolph 1994; Diaz-Uriarte and Garland 1996; Martins 1996; Oakley and Cunningham 2000).

A widely used approach for testing correlation between traits on a phylogenetic tree is the independent contrasts method of Felsenstein (1985) which extracts independent quantities from the tip values of the traits in order to estimate their correlation. The rationale behind this approach is that if two traits follow two correlated Brownian motions then their matched independent contrasts are realizations of independent and identically distributed pairs of Gaussian random variables correlated with the same correlation as the Brownian motions. Therefore, testing for correlation through independent contrasts is perfectly founded if one assumes that the two traits to compare follow Brownian motions along the phylogenetic tree, thus a neutral evolution for both of them (Felsenstein 1988). A strongly related method, called Phylogenetic Generalized Least Squares (PGLS, Grafen 1989) addresses the same question in an equivalent way. We extend and give formal proofs of results about the equivalence between independent contrasts and PGLS methods for regression analysis stated in Garland and Ives (2000); Rohlf (2001); Blomberg et al. (2012).

Both independent contrasts and PGLS approaches make the strong assumption that evolution of the considered traits is neutral. There are many situations where this assumption is not granted, for instance when evolution is driven by ecological pressures. Arnold and Moncrieff (1994) showed that several lizard species developed the same discrete traits in approximately the same order when adapting to a same environmental condition. In the same way, quantitative traits may evolves toward optima or following a general trend. This type of non-neutral evolution, referred to as adaptative and directional evolution, cannot be modeled by using standard Brownian processes. Adaptation, i.e., evolution toward optima, is generally modeled with Ornstein-Uhlenbeck processes or more complex models (Cressler et al. 2015). For instance, Hansen et al. (2008) consider traits evolving according Ornstein-Uhlenbeck processes with optima which themselves evolve following a Brownian process. Directional evolution is generally modeled with arithmetic Brownian motions, i.e., Brownian process with linear trends. Note that after a change in the optimum during the adaptation process, traits under selection may look under directional evolution until getting close to their new optima. In this work, we focus on the directional evolution case. More precisely, we address the question of how to detect correlation between two traits when at least one of them is under directional evolution. Among the numerous examples of evolutionary tendencies in the evolution of phenotypic or genetic traits, Cope’s rule, which predicts that the body size of species tends to increase over evolutionary time, has gained considerable empirical support (Kingsolver and Pfennig 2004; Van Valkenburgh et al. 2004; Hone and Benton 2005; Hone et al. 2005; Hunt and Roy 2006; Bokma et al. 2016). Beside the Cope’s rule, there are strong evidence that traits of some clades have evolved following trends at certain periods. For instance, there are evidence of increase in the Mysticetes body size (Slater et al. 2010) and in the microsatellite size in Maize (Vigouroux et al. 2003). Last, insular dwarfism and gigantism phenomena provide numerous examples of directional evolution (Lomolino 2005). We emphasize that an evolutionary tendency in increasing or decreasing the body size of species implies that most of their morphological measures follow the same trend, and would be systematically tested as significantly correlated (Yule 1926; Entorf 1997; Deng 2015).

In order to study the correlation between traits under directional evolution, we first determine the probability distribution of the independent contrasts of a trait which evolves following a Brownian motion with a linear trend. The form of these distributions shows that testing for correlation between two traits under directional evolution through independent contrasts makes no sense. We propose an alternative approach based on a multiple regression which includes time as explanatory variable in order to correct the bias due to a linear trend. We show that our approach is equivalent to performing a PGLS analysis by adding the tip times as covariable in the regression.

A previous approach to correct the trend effect on the independent contrasts and to test for correlation between traits under directional evolution was proposed in Elliott (2015). Its general idea is to “center” the independent contrasts with regard to the trend (this method is detailed below). A thorough study of the Elliott’s (2015) approach shows that it does not satisfy the regression assumptions. Note that if the phylogenetic tree supporting the evolution of the traits is ultrametric, our new method and that of Elliott (2015) are both equivalent to the independent contrasts method.

We simulated evolution of correlated and uncorrelated quantitative traits with and without trend on hominin phylogenetic tree in order to assess and to compare our new correlation test and three previous ones, namely the standard correlation test on the tips values of the traits, the correlation between independent contrasts of Felsenstein (1985) and that between the directional contrasts proposed by Elliott (2015). Simulation results shows that the new test is the most accurate as soon as one of the traits is under directional evolution. Despite its statistical flaws, the approach suggested by Elliott (2015) performs almost as well as our approach.

Last, the approach was applied on a real dataset in order to test for correlation between the logarithms of hominin body mass and cranial capacity, among which the logarithm of cranial capacity shows a significant positive trend. Our test concluded to a significant correlation between the logarithms of these two traits.

R-scripts implementing the correlation tests and the simulations performed for this work are available at https://github.com/gilles-didier/Correlation.

The rest of the paper is organized as follows. The independent contrasts and their distributions when traits follow Brownian motions with linear trends are presented in Section 2. Section 3 recalls three previous correlation tests on phylogenetic data and introduces a new one based on a multiple regression between independent contrasts which includes time as explanatory variable. The four correlation tests are assessed and compared on simulated data in Section 4. Last, our test is applied to check correlation between hominin cranial capacity and body mass in Section 5.

## 2 Independent contrasts

### 2.1 Phylogenetic trees - Notations

In all what follows, we assume that the evolutionary history of the species is known and given as a rooted binary phylogenetic tree *𝒯* with branch lengths. Our typical tree *𝒯* contains 2*n* + 1 nodes among which *n* are internal nodes. According to the Felsenstein’s (1985) convention, the nodes are indexed in the following way:

- index 0 for the root,
- indices 1 to *n* − 1 for the other internal nodes,
- indices *n* to 2*n* for the tips.

For all nodes *k*, we put

- *r*(*k*) and *l*(*k*) for the two direct descendants of *k*, if *k* is an internal node,
- *a*(*k*) for the direct ancestor of *k*, if *k* is not the root,
- *v*_*k*_ for the length of the branch ending at *k*,
- *t*_*k*_ for the (absolute) time of *k*, which is the sum of the branch lengths of the path relying the root to *k* (both included), and
- *z*_*k*_ for the value of the trait at node *k*, which is defined only if *k* is a tip.

### 2.2 Felsenstein’s (1973) algorithm

Independent contrast were introduced in Felsenstein (1973) in order to compute the probability of the tip values of a quantitative trait evolving on a phylogenetic tree under the assumption that this trait follows a Brownian motion. This method extracts a series of realizations of independent Gaussian variables from the tip values by iteratively considering differences between terminal sister taxa and by replacing them with a single terminal taxa, while modifying the length of the branch that it ends and associating it with an artificial trait value computed from those of the two terminal sister taxa. Namely, the method recursively computes a new branch length 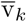 and an “artificial” trait value 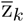 for all nodes *k* of *𝒯* in the following way.

#### Definition 1

(Felsenstein 1973). *Let 𝒯 be a phylogeny and* (*z*_*k*_)_*n*≤*k*≤2*n*_ *be the values of a quantitative trait only known at the tips of 𝒯*. *Under the notations of Section 2.1 and for all nodes k of 𝒯, the quantities* 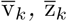 *are recursively defined as*

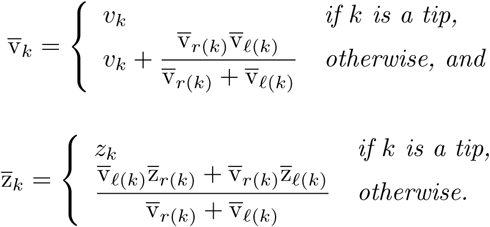

*For all internal nodes k of 𝒯, the (standardized) independent contrast u*_*k*_ *is then defined as*

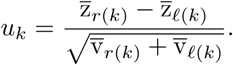

Contrasts can be computed thanks to the `pic` function in the ape `R-package` (Paradis et al. 2004).

### 2.3 Independent contrasts as random variables

By construction, under an evolutionary model for quantitative traits, the tips values (*z*_*k*_)_*n*≤*k*≤2*n*_ are realizations of random variables *Z*_*n*_, …, *Z*_2*n*_. In this context, the artificial trait values 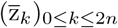 computed during the independent contrasts procedure are also realizations of random variables which can be expressed from *Z*_*n*_, …, *Z*_2*n*_ and so are the independent contrasts.

#### Definition 2.

*Let* (*Z*_*k*_)_*n*≤*k*≤2*n*_ *be the random variables associated to the tip values of a trait. For all nodes k of 𝒯, the random variable* 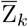 *is defined as*

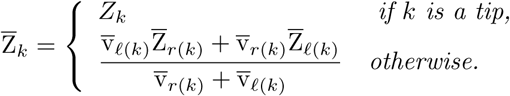

*And for all internal nodes k of 𝒯, the random variable U*_*k*_ *is defined as*

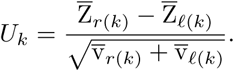

In plain English, for all nodes *k*, the random variable 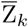 corresponds to the value associated to *k* by the computation presented in Definition 1. In the same way, for all internal nodes *k*, the contrast *u*_*k*_ is a realization of the random variable *U*_*k*_.

As evolutionary models of traits, we shall consider below either the Brownian Motion (BM) model or the Brownian Motion with linear trend, also known as the Arithmetic Brownian Motion (ABM) model. Namely, the ABM model with parameters (*x*_0_, *µ, σ*^2^), i.e., initial value *x*_0_, trend *µ* and variance *σ*^2^, is the process (*X*_*t*_)_*t*>0_ defined as

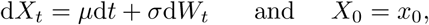

where (*W*_*t*_)_*t*>0_ is the Wiener process (Grimmett and Stirzaker 2001). For all times *t* and *s*, the increments 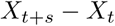 are independent Gaussian random variables with law 𝒩 (*µs, σ*^2^*s*).

Since a BM model is nothing but an ABM model with trend parameter *µ* = 0, we shall write results and properties in the ABM case only. Basically, any property or result granted for ABM also holds for BM.

The ABM process with parameters (*x*_0_, *µ, σ*^2^) running on the phylogenetic tree *𝒯* starts at the root of 𝒯 with the value *x*_0_, then evolves independently on each branch of 𝒯 by splitting at each internal node into two independent and identical processes both starting from the value of the process at this node and eventually ends at the tips of 𝒯. It allows to model traits under directional evolution, e.g., following the Cope’s rule (Kingsolver and Pfennig 2004; Van Valkenburgh et al. 2004; Hone and Benton 2005).

#### Theorem 3.

*Let* (*Z*_*k*_)_*n*≤*k*≤2*n*_ *be the random variables associated to the tip values of a trait following the ABM model with parameters* (*x*_0_, *µ, σ*^2^) *on 𝒯*. *For all nodes k of 𝒯, the random variable* 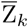 *can be written as*

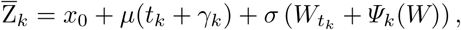

*where*

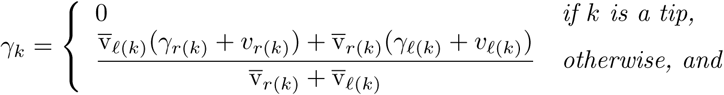

*and Ψ*_*k*_(*W*) *is a linear combination of increments of the form 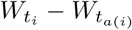, recursively defined as*

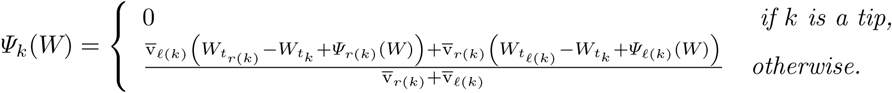

*For all internal nodes k of 𝒯, the random variable U*_*k*_ *can be written as*

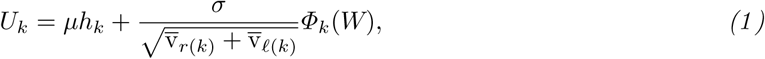

*where*

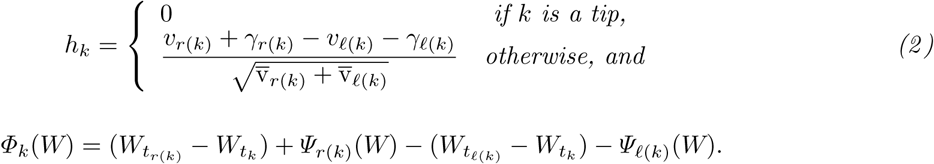

*Random variables Ψ*_*k*_(*W*) *and Φ*_*k*_(*W*) *have Gaussian distributions* 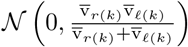 *and* 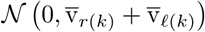 *respectively*.

*Proof*. Appendix A.

**□**

#### Corollary 4.

*Under the ABM model with parameters* (*x*_0_, *µ, σ*^2^), *for all internal nodes k of 𝒯, the independent contrast random variables U*_*k*_ *are independent from one another and Gaussian distributed with*

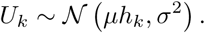

*Proof*. The proof of the independence of the independent contrasts under the ABM model follows the same arguments as the proof of the same property in the BM case provided in Felsenstein (1973). The form of the distribution of the independent contrasts is a direct consequence of Theorem 3. □

#### Proposition 5.

*Let 𝒯 be an ultrametric tree and T be the length of the path from the root to the tips. For all nodes k of 𝒯, we have that γ*_*k*_ = *T* − *t*_*k*_ *and h*_*k*_ = 0.

*Proof*. Appendix B. □

### 2.4 Correlation and independent contrasts

Independent contrasts are mainly used in order to test for correlation between two quantitative traits known only at the tips of a phylogenetic tree (Felsenstein 1985). The rationale behind this approach is that if two quantitative traits follow two correlated Brownian motions then their standardized independent contrasts are realizations of independent and identically distributed pairs of Gaussian random variables with the same correlation as the correlated Brownian motions. The transpose of a matrix or a vector *D* is noted *D*′

#### Definition 6.

*Let* 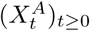 *and* 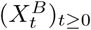 *two ABM models with parameters* 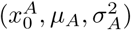 *and* 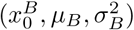 *respectively and let* 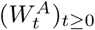 *and* 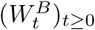 *be two Wiener processes such that*

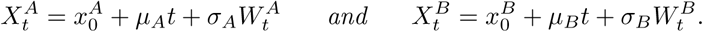

*The processes* 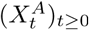 *and* 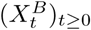 *are correlated with correlation coefficient ρ if for all t* ≥ 0, *the random vector* 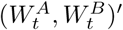 *has covariance matrix t*∑ *where*

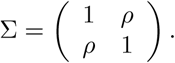

Any pair of Wiener processes 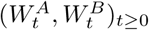 with correlation matrix ∑ can be obtained from two independent Wiener processes 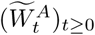 and 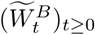 by applying the Cholesky decomposition on their covariance matrix ∑ (Gupta 2013). Namely, by decomposing ∑ as ∑ = *L L*′where

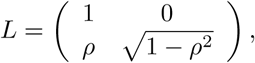

the random vector 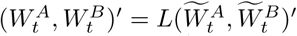 has covariance matrix ∑.

In sum, two ABM processes 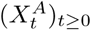 and 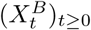 with parameters 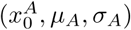 and 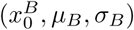 are correlated with correlation coefficient *ρ* if and only if they can be written as

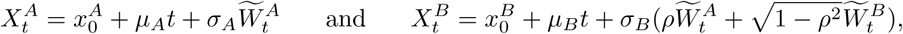

where 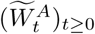 and 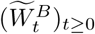 are two independent Wiener processes.

#### Theorem 7.

*Let* 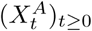 *and* 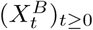 *be two ABM processes running on 𝒯 with parameters* 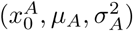 *and* 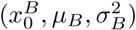 *respectively and* 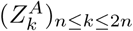 *and* 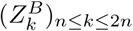 *be the random variables associated to the values of these processes at the tips of 𝒯. The processes* 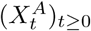 *and* 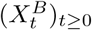 *are correlated with correlation coefficient ρ if and only if for all internal nodes k of 𝒯, the independent contrast random variables* 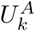 *and* 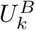 *computed from* 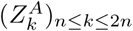 *and* 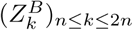 *are correlated with correlation coefficient ρ*.

*Proof*. Let us assume that the ABM processes 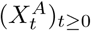 and 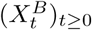 of parameters 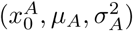 and 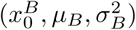 respectively are correlated with correlation coefficient *ρ*. It is equivalent to say that there exist two independent Wiener processes 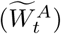 and 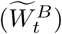 such that

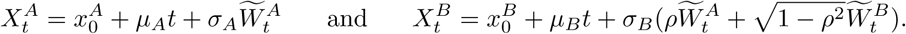

By applying Equation 1 of Theorem 3 to 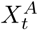 and 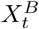, we then have that for all internal nodes *k*, the independent contrast random variables 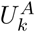 and 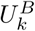 can be written as

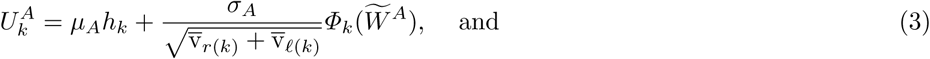

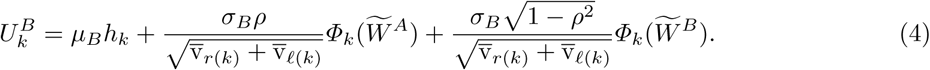

Since Theorem 3 ensures that var 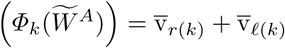, it follows that

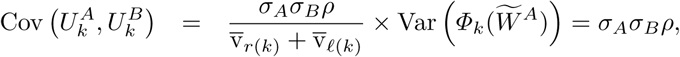

thus 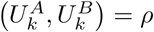 for all internal nodes *k* of *𝒯*, which ends the proof. □

## 3 Correlation tests

In practical situations, evaluating or testing for correlation between two random variables is performed by considering a series of a certain number *n* of independent realizations of this pair of random variables. Let us remark that sampling independently *n* times a pair of random variables is equivalent to draw a joint sample of *n* independent and identically distributed pairs of random variables with the same correlation between the random variables of all the pairs. Standard regression analysis can thus be applied in this last case.

We emphasize the fact that performing a correlation test on a joint sample of *n* pairs of random variables which are not identically distributed makes absolutely no sense, even if each pair has the same correlation. Moreover, if we do not have independence between the pairs of random variables, then assumptions required by correlation tests are violated.

Linear regression is a usual tool for studying the association between two variables. Statistic analysis of regression requires additional assumptions, which are referred to as the *key assumptions* in Fox (2015), namely the constancy of the error variance, the Gaussianity of the errors, their null mean, and their independence. In the multiple linear regression, testing for correlation between the response variable and one of the regressors is performed by testing the nullity of the corresponding regression coefficient. Under the key assumptions, this test of nullity is based on the fact that, by assuming that the coefficient is null, the ratio of the ordinary least squares estimate of this coefficient to its standard deviation follows a Student distribution with a number of degrees of freedom equal to the difference between the number of samples and the number of regressors (including the intercept if there is one, Fox 2015). In the case of the simple linear regression with intercept, testing the nullity of the regression coefficient corresponding to the slope is equivalent to performing a Pearson’s correlation test between the response and the explanatory variables (Kendall and Stuart 1961, p985).

Let *A* and *B* be two traits, 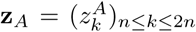 and 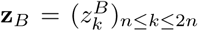 be their tip-value vectors (of dimension *n* + 1) and 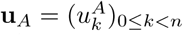 and 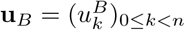 be the corresponding independent contrast vectors (of dimension *n*) and 1 be for the vector with all entries equal to 1 (its dimension depending on the context). Below, we shall present several ways of testing for correlation between *A* and *B*.

### 3.1 Standard regression (SR)

The most basic way to test for correlation between two traits *A* and *B* is to consider the standard Pearson’s correlation test obtained from the linear equation:

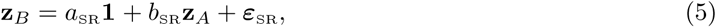

which will be referred to as the SR method. In the SR method, the Pearson’s correlation test between traits *A* and *B* amounts to testing for the nullity of the coefficient *b*_SR_.

Let

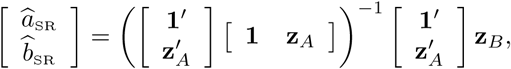

be the vector of ordinary least square estimates of the coefficients *a*_SR_ and *b*_SR_. The variance estimate of *b*_SR_ is 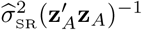 where

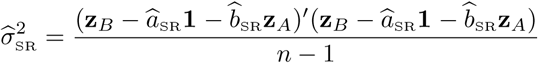

is the residual variance estimate. Under the key assumptions of the regression model, testing the nullity of the coefficient *b*_SR_ is performed thanks to the fact that if *b*_SR_ = 0 then the ratio of the coefficient estimate 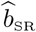 to its standard error 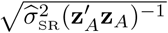 follows a Student distribution with *n* − 1 degrees of freedom. Unfortunately the key assumptions of the regression model are not granted here since entries of the error vector ***ε***_SR_ are not sampled from independent and identically distributed random variables because of the evolutionary relationships between the species involved (Harvey and Pagel 1991). Though one expects correlation tests with the SR method to be inaccurate in a phylogenetic context (except if *𝒯* is a star tree), the SR method is included in the study in order to be used as a basis of comparison.

### 3.2 Independent contrasts (IC)

The usual way to cope with the evolutionary dependency between the tip values of the traits is to consider their independent contrasts (Felsenstein 1985), referred to as the IC method below, which is based on the regression through origin between the independent contrasts according to the equation:

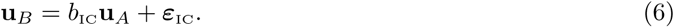

The IC method is widely used and has been assessed in several works (Grafen 1989; Martins and Garland 1991; Pagel 1993; Garland and Adolph 1994; Martins 1996; Diaz-Uriarte and Garland 1996). In the IC approach, testing correlation between traits *A* and *B* amounts to testing for the nullity of the coefficient *b*_IC_ in the regression through the origin. The ordinary least square estimate 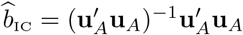 of *b*_IC_ has variance 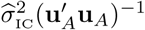 where

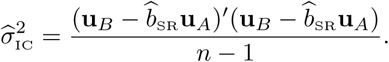

Under the key assumptions of the regression analysis, if *b*_IC_ = 0 then the ratio of 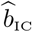to its standard deviation follows a Student distribution with *n* − 1 degrees of freedom.

Testing for the nullity of *b*_IC_ by using this property is theoretically founded if one assumes that both traits *A* and *B* evolve following a BM model since Corollary 4 and Theorem 7 ensures that their respective independent contrasts are realizations of independent and identically distributed centered Gaussian random variables 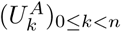 and 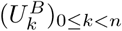 with the same correlation as the two traits. Putting 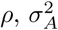 and 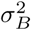 for this correlation and the variances of 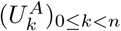 and 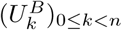, respectively, and considering a Cholesky decomposition of their covariance matrix, we get that the random contrasts 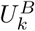 can be written as

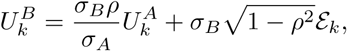

where the terms 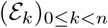 are independent centered standard Gaussian variables. This shows that the key assumptions of the regression analysis are well granted. The correlation between the independent contrast random variables 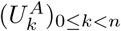 and 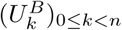, thus between the traits *A* and *B*, can be assessed by regression through origin between the contrast series 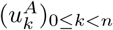 and 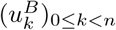.

In the case where at least one of the two traits to compare follows an ABM model with a non-zero trend, Theorem 7 still ensures that their respective independent contrasts are correlated with the same correlation as the traits. Unfortunately, since from Corollary 4 their independent contrasts are no longer identically distributed but depend on *k* (we have 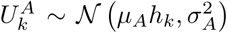 and 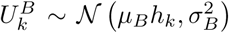 with *h*_*k*_ ≠ 0 in the general case), it makes no sense to test for their correlation through a joint sample.

Figure 1 illustrates how directional trends may make independent contrasts computed from two uncorrelated traits look strongly correlated. This was expected since spurious correlations due to a common dependency on a third factor is a classical phenomenon (Yule 1926; Entorf 1997; Deng 2015).

**Figure 1:**
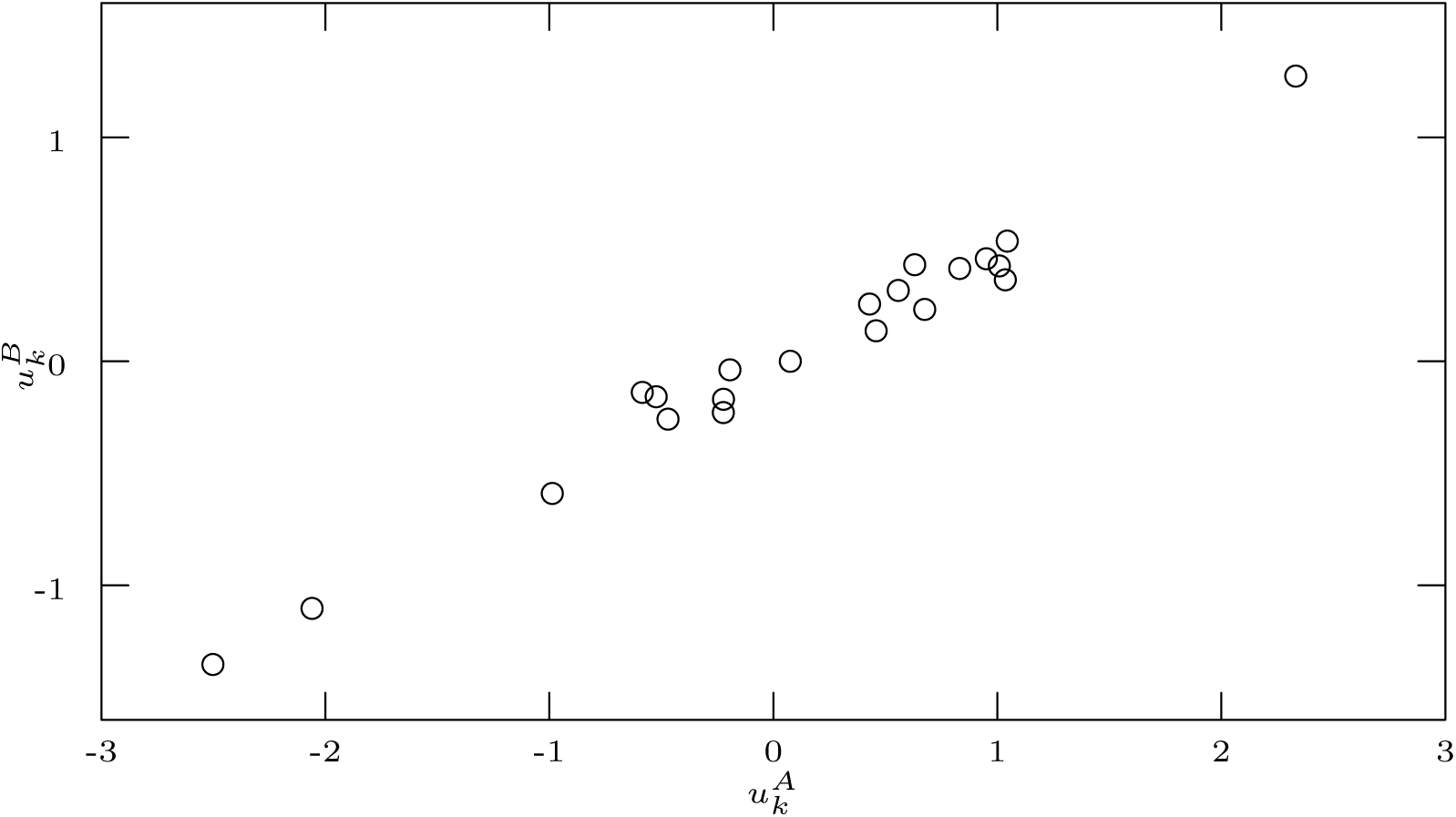
Independent contrasts of two traits *A* and *B* simulated on the tree of Figure 2 under two uncorrelated ABM models with trends *µ*_*A*_ = 0.5 and *µ*_*B*_ = 0.2 respectively.

### 3.3 Directional contrasts (DC)

A first approach to correct the independent contrasts when at least one of the two traits to compare evolves with a linear trend was proposed in Elliott (2015). The general idea of Elliott’s (2015) approach is to center the independent contrasts in order to make them identically distributed. To this end, Elliott (2015) defined the *β*-directional contrasts *d*_*k*_(*β*). The formal definition of the *β*-directional contrasts is recalled in Appendix D in which we show that, for all internal nodes *k*, the *β*-directional contrasts *d*_*k*_(*β*) of Elliott (2015) are equal to the *β*-centered contrasts *c*_*k*_(*β*) defined as:

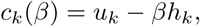

which have a direct interpretation with regards to the formalism of Section 2.

Let us first remark that under an ABM model with trend *µ*, Theorem 3 ensures that the *µ*-centered/*µ*- directional contrasts are well independent and identically distributed thus could be used in correlation tests. Unfortunately, obtaining these corrected contrasts requires to have the trend parameter *µ*. More exactly, for testing correlation between two traits *A* and *B*, their respective trends *µ*_*A*_ and *µ*_*B*_ have to be known, but they are *a priori* unknown in practical situations. Elliott (2015) proposed to use instead their estimates 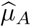 and 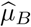 (Equation C2) and to consider the correlation between the (estimated) directional contrasts 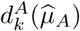 and 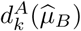, where 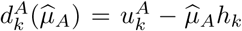 and 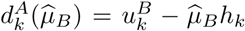 for all internal *k* nodes of. 𝒯

Regression between estimated directional contrasts will be referred to as the DC method. Namely, by putting **d**_*A*_ and **d**_*B*_ for the vectors of directional contrasts 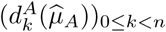 and 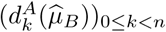, it is based on the following linear equation:

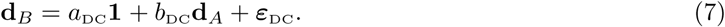

Let

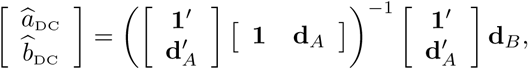

be the vector of ordinary least square estimates of the coefficients *a*_DC_ and *b*_DC_ and

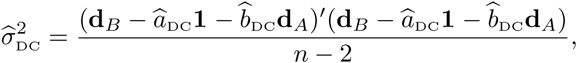

be the residual variance (vectors **d**_*A*_ and **d**_*B*_ have dimension *n*). Testing for correlation between traits *A* and *B* amounts to testing for the nullity of the parameter *b*_DC_, which, under the key assumptions of the regression model, is performed thanks to the fact that, by assuming that *b*_DC_ = 0, the ratio of the coefficient estimate 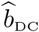 to its standard error 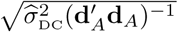 follows a Student distribution with *n* − 2 degrees of freedom.

Unfortunately, since the trends 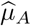 and 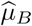 are estimated from independent contrasts, the estimated directional contrasts are neither independent, nor identically distributed under ABM models (Appendix D). Applying standard correlation tests on estimated directional contrasts is not founded from a statistical point view.

### 3.4 Multiple Regression (MR)

If traits *A* and *B* follows two ABM models 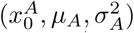 and 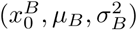 then, for all internal nodes *k* of *𝒯*, both independent contrast random variables 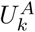 and 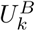 depend on the same explanatory variable *h*_*k*_ (Equation 1). As shown in Figure 1, this dependence on a common factor is likely to cause a systematic correlation between the random variables 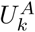 and 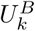. Neutralizing this spurious correlation requires to include the common explanatory variable *h*_*k*_ in the regression (Yule 1926; Deng 2015). By combining Equations 3 and 4, we get that

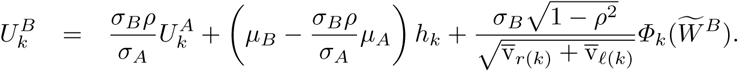

This suggests to consider the multiple regression through origin between contrasts by including *h*_*k*_ as co-variable, i.e., to consider the equation:

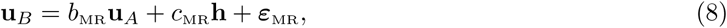

which will be referred to as the MR method. The multiple regression procedure is statistically sound here since the entries of the error vector are sampled from random variables 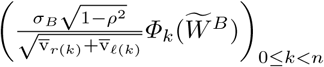 which are independent and Gaussian distributed with mean zero and constant variance 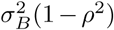 under the current assumptions. The vector of ordinary least square estimates of *b*_MR_ and *c*_MR_ is

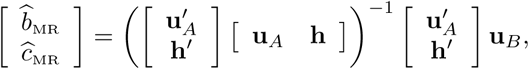

and the variance of the estimator of *b*_MR_ is 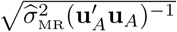 where

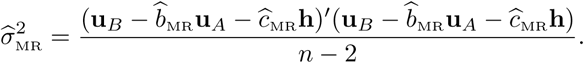

Equations 3 and 4 show that if traits *A* and *B* follows two ABM models 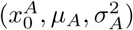 and 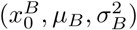 with correlation *ρ*, the random contrasts 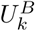 can be written as

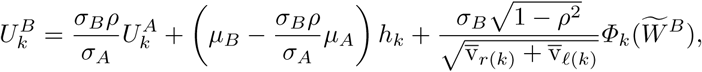

where the random error variables 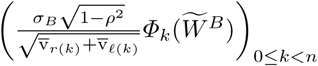 are independent and Gaussian distributed with mean zero and constant variance 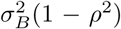. The key assumptions of the regression analysis are thus granted if traits *A* and *B* follow two ABM models. Testing for correlation between traits *A* and *B* can be performed by testing for the nullity of coefficient *b*_MR_ thanks to the fact that if *b*_MR_ = 0 then the ratio of the coefficient estimate 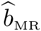 to its standard deviation follows a Student distribution with *n* − 2 degrees of freedom.

It is worth pointing out that the MR test does not require to estimate neither trend *µ*_*A*_ nor trend *µ*_*B*_.

### 3.5 Relation with PGLS method

PGLS method was introduced in Grafen (1989) and further studied in Martins and Garland (1991); Pagel (1997); Martins and Hansen (1997). It is a generalized least squares method specifically designed to take into account the phylogenetic dependency of the regression errors. This dependency relies on evolutionary assumptions. In particular under the BM model, the dependency structure is exactly the same as for the IC method which is based on the same model. Namely, the covariance matrix of the tip random variables of 𝒯has the form *σ*^2^∑ where *σ*^2^ is the variance of the Brownian model and ∑ is the matrix indexed on the tips of 𝒯 such that for all pairs of tips (*i, j*), the entry (*i, j*) is the total time between the root of 𝒯 and the most recent common ancestor of *i* and *j*. The PGLS approach is based on the following linear equation

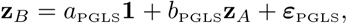

which looks the same as that of the SR method but the error vector *ε*_PGLS_ is now assumed to be sampled from a centered Gaussian vector with covariance matrix proportional to ∑. The vector of general least square estimates of the coefficients *a*_PGLS_ and *b*_PGLS_ is

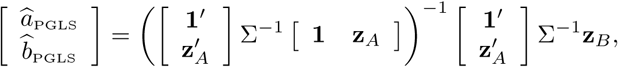

and the residual variance estimate is

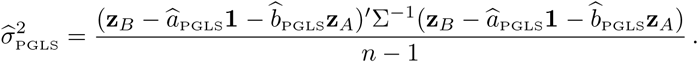

Under the PGLS assumptions, testing the nullity of the coefficient *b*_PGLS_ is performed thanks to the fact that, by assuming that *b*_PGLS_ = 0, the ratio of the coefficient estimate 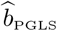 to its standard error follows a Student distribution with *n* − 1 degrees of freedom.

The strong relation between PGLS and IC approaches is well known (Garland and Ives 2000; Rohlf 2001). Blomberg et al. (2012) provided a formal proof that, in the simple regression case, least square estimates of the regression coefficient of the explanatory variable is exactly the same with the IC as with the PGLS methods, namely that 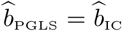. In Appendix E, we prove the same result in the multiple regression case and show that the variance of the least square estimates of the coefficients are also the same with the IC and the PGLS methods.

#### Theorem 8.

*Let* **z**_0_, **z**_1_,…**z**_*p*_ *be the tip value vectors of traits or covariables (e*.*g*., *tip times, environmental variables* …*) and* **u**_0_, **u**_1_,…**u**_*p*_ *be the corresponding independent contrast vectors*, 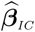 *be the vector of the ordinary least square coefficient estimates from the linear equation though origin*

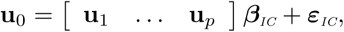

*and let* 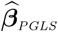 *be the vector of the generalized least square coefficient estimates from the linear equation with intercept*

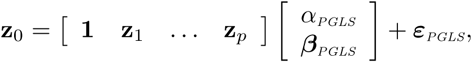

*where the error vector ε*_*PGLS*_ *is assumed to be a realization of a centered Gaussian vector with covariance matrix proportional to* ∑ *(the covariance matrix associated to the tree 𝒯). The vectors* 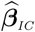 *and* 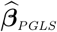 *as well as covariance matrices of the corresponding least squares estimators are equal. Moreover, the degrees of freedom involved in the nullity tests of their coefficients are both equal to n* − *p with the two approaches*.

*Proof*. Appendix E. □

Theorem 8 directly implies that testing for correlation between two traits with IC and PGLS is completely equivalent, even by considering others traits or covariables.

In order to show the relation between the MR and the PGLS approaches, let us add the tip times as an explanatory covariable in the PGLS regression, i.e., let us consider the linear equation

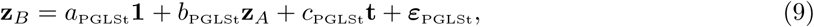

where **t** = (*t*_*k*_)_*n*≤*k*≤2*n*_ is the tip time vector. The vector of general least square estimates of the coefficients *a*_^PGLSt^_, *b*_^PGLSt^_ and *c*_PGLSt_is

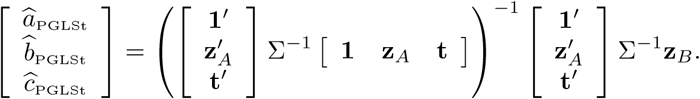

Testing for the nullity of *b*_PGLSt_ is performed by considering the ratio of 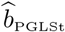 to its standard deviation in the Student distribution with *n* − 2 degrees of freedom.

We prove in Appendix E that the variables (*h*_*k*_)_0≤*k<n*_ are the independent contrasts of the tip times. Theorem 8 then implies that both the least square estimates of the regression coefficient associated to trait *A* and their variances are exactly the same with the MR method and with the PGLSt method.

In short, the IC and PGLS (resp. the MR and PGLSt) methods are interchangeable to test for correlation between two continuous traits under neutral or directional evolution.

An important point is that computations of the PGLS method require to inverse the covariance matrix accounting for the phylogenetic dependencies, which has cubic time complexity with respect to the size of the tree, whereas the IC and MR methods takes advantage of the tree structure of the phylogenetic dependencies in order to perform the same computations in linear time.

### 3.6 Ultrametric trees

Proposition 5 states that if 𝒯 is ultrametric then *h*_*k*_ = 0 for all internal nodes *k* of 𝒯. This implies that Equations 6 and 8 turn out to be exactly the same in this case. In other words, on an ultrametric phylogenetic tree, the IC and MR methods are totally equivalent to test for correlation. Moreover, since in an ultrametric tree, the maximum likelihood estimator of the trend returns always 0 (again because *h*_*k*_ = 0 for all internal nodes *k* of 𝒯, cf Equation C2), Equation 7 is the same as Equations 6 and 8. In sum, the IC, DC and MR methods are equivalent on ultrametric trees. Using the IC method is statistically founded here since independent contrasts satisfy the requirements of correlations tests in the ultrametric case.

Note that non-ultrametric phylogenetic trees arise in several situations. In particular, phylogenetic trees containing fossil taxa (with or without extant taxa) are not ultrametric (e.g., Laurin 2004; Heim et al. 2015). The ultrametric character relies on the evolutionary model used to infer the trees. For instance, the speciational model which somehow assumes a same unitary branch length all along the tree generally provides non-ultrametric trees (Knouft and Page 2003; Moen 2006; Laurin et al. 2012). Measuring branch lengths in terms of genetic changes (Moen 2006) or scaling branch lengths by their own evolution rates in heterogeneous models (Baker et al. 2015, 2016) instead of considering their geological ages also lead to non-ultrametric trees even if all the taxa are extant.

## 4 Simulation study

In this section, we shall assess and compare the four correlation tests presented in Section 3:

- SR: standard regression of tips values (Equation 5),
- IC: regression through origin of independent contrasts (Equation 6),
- DC: regression of directional contrasts (Equation 7),
- MR: multiple regression through origin of independent contrasts with *h*_*k*_ as co-variable (Equation 8).

### 4.1 Simulation and evaluation protocol

We simulated the evolution of two quantitative traits *A* and *B* under various conditions, i.e., under BM and ABM models with several sets of parameters and several levels of correlation between *A* and *B*. The simulated evolution runs on the hominin phylogenetic tree displayed in Figure 2 (Dembo et al. 2015).

**Figure 2:**
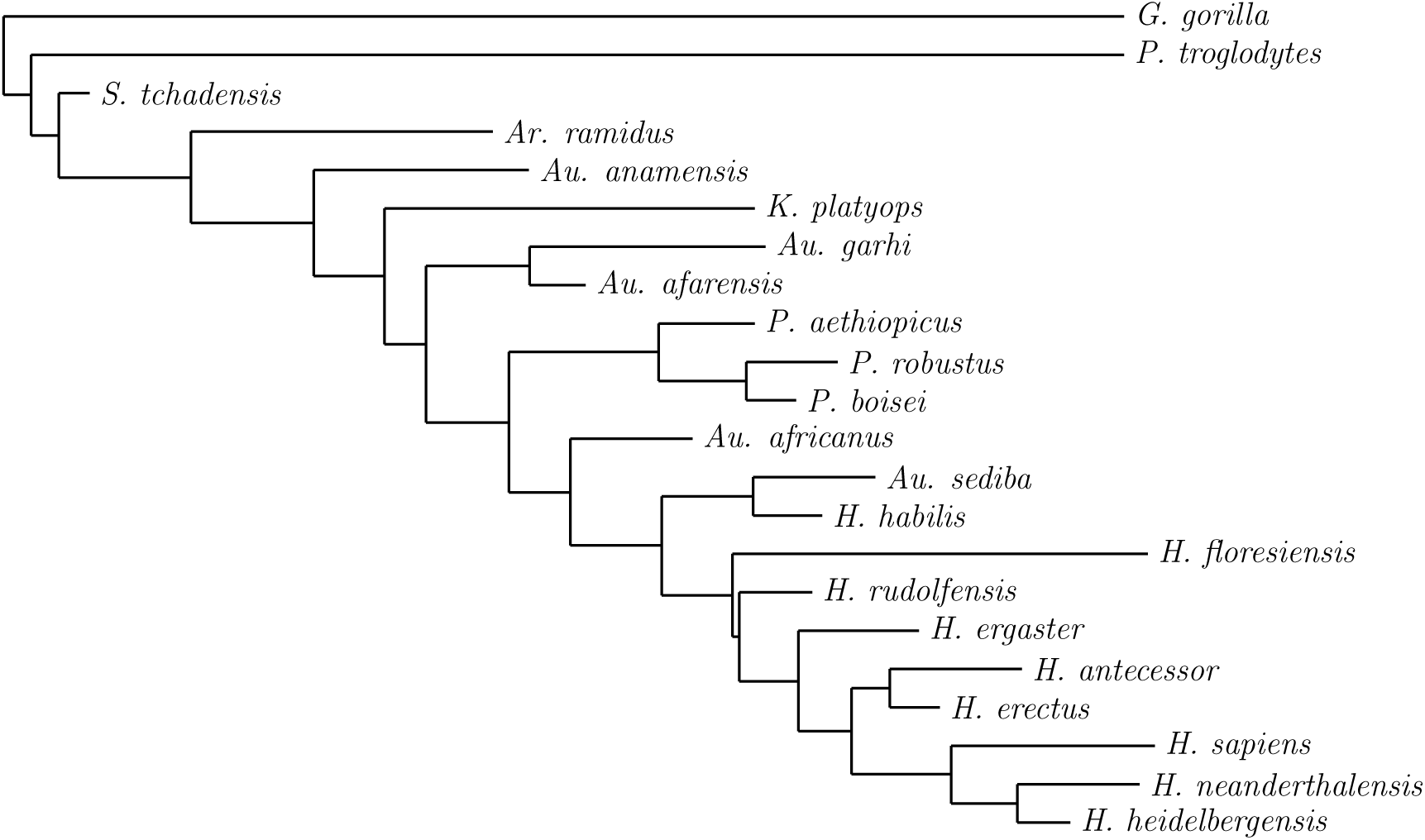
Hominin phylogeny (Dembo et al. 2015).

Although the ABM model has three parameters (*x*_0_, *µ, σ*^2^), we only vary the trend parameter *µ* in the simulations. The parameter *x*_0_ just translates the whole evolution process, which has no effect on the correlation of a trait with another. Multiplying both the trend and the standard deviation of an ABM model with a constant just results in multiplying the values of the ABM process with same constant (i.e., what actually matters is the ratio of the trend to the standard deviation).

The four correlation tests were next assessed in terms of type I error, i.e., with regard to their ability to not falsely reject the null hypothesis, the null hypothesis being that the traits are uncorrelated, in the case where the traits to compare are actually uncorrelated. We also displays ROC plots of the tests for summarizing their ability to distinguish between correlated and uncorrelated traits (Zhou et al. 2011). Plots of type I error were obtained by simulating uncorrelated traits and by plotting the proportion of simulations for which the null hypothesis was rejected *versus* the level of risk (each test associates to a simulation, a level of risk between 0 and 1, accounting for the chance that this simulation satisfies the null hypothesis). ROC curves were obtained by simulating both negative (i.e., uncorrelated) population and positive (i.e., correlated) population and by plotting for all levels of risk the proportion of true negatives *versus* the proportion of false positives detected by each test. We simulated 50 000 evolutions of correlated and uncorrelated pairs of traits for each plot.

### 4.2 Correlation tests between two traits under neutral evolution

We first simulated two traits evolving under neutral evolution (i.e., under the BM model) with correlation levels 0 and 0.5 (results obtained with correlation 0.7 are provided in the supplementary information).

Figure 3-Left displays the proportion of type I error at all level of rejection *α* obtained from 50,000 simulations of two uncorrelated traits under the BM model with variance 0.09. We do observe that both the IC and the MR methods are perfect in the sense that they both rejected the null hypothesis at the exact level *α* required (both plots of IC and MR completely overlap with the diagonal in Figure 3-Left). The DC method is close to perfect but tends to reject the null hypothesis a little bit more than it should do. Last, as expected, the worst performance comes from the SR method.

**Figure 3:**
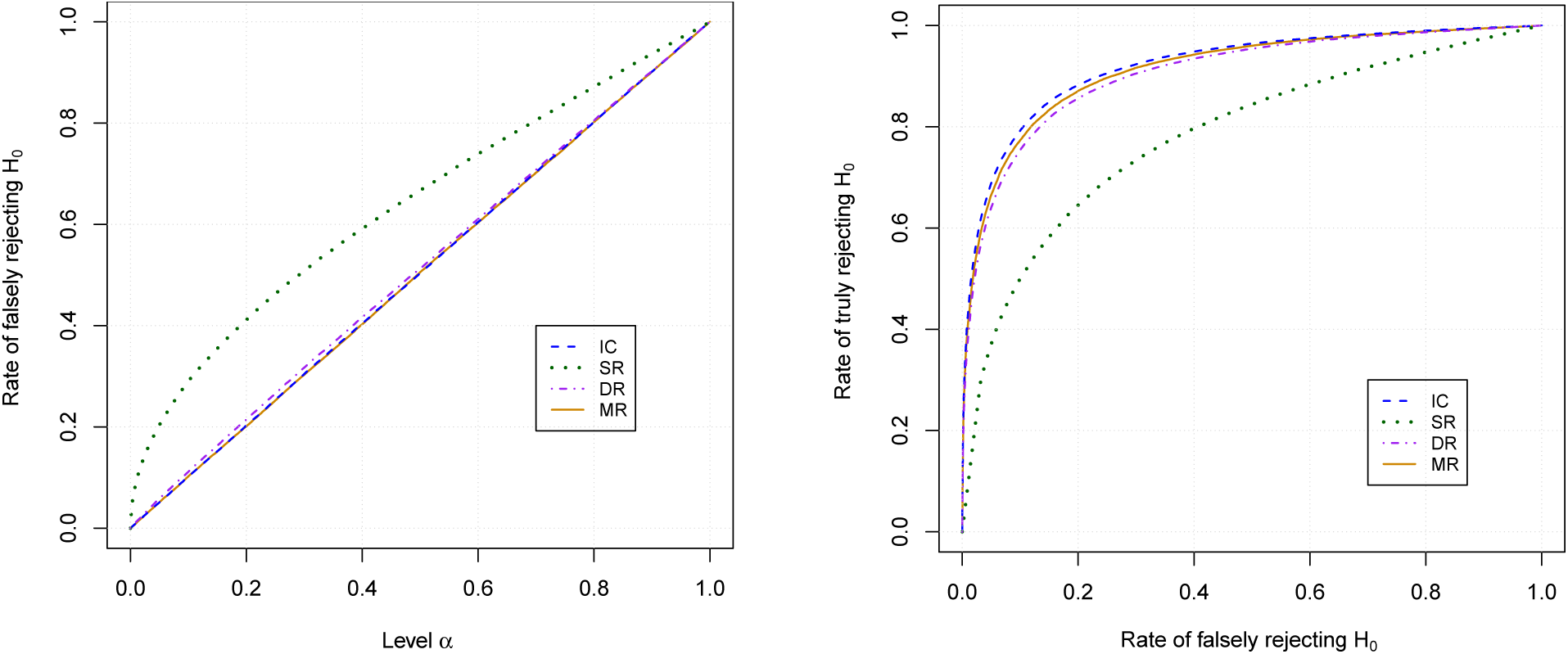
Left: Rates of false rejection of the null hypothesis at level *α vs α* when both traits *A* and *B* follow the BM model with variance 0.09. Right: ROC plots of the correlation tests obtained from two simulated traits under the BM model with variance 0.09. Negative population is simulated with uncorrelated traits and positive population with traits correlated with correlation 0.5.

The *p*-values obtained from testing for the simulations are expected to be uniformly distributed with support [0, 1]. We used Kolmogorov-Smirnov (K-S) test in order to check this point. We observed that *p*-values obtained when testing for the correlation of independent traits follow an uniform distribution both for the IC method (Kolmogorov-Smirnov test, *p*-value=0.537) and for the MR method (K-S test, *p*-value=0.832). This is the case neither for the DC nor for the SR methods (K-S test, *p*-values smaller than 10^*-*8^). In sum, under the BM model, only the IC and the MR methods have the behavior expected from a statistical test.

The ROC plots of the tests with a positive population simulated under the same BM model, but with a correlation 0.5 between the traits, are displayed Figure 3-Right. It shows that under the BM model, the most accurate test is IC but MR and DC tests have close performances. As expected the less accurate test is SR.

### 4.3 Correlation tests between a trait under neutral evolution and a trait under directional evolution

We consider here the mixed situation where one of the traits follows a neutral evolution, here simulated under the BM model with variance 0.09 and the other one follows a directional evolution, here simulated under the ABM model with trend 0.5 and variance 0.09.

Figure 4-Left shows that the behavior of the type I error with regard to the level of rejection *α* is essentially the same as in the case of two traits under neutral evolution for all the methods. Moreover, *p*-values obtained here still follow an uniform distribution with support [0, 1] both for the IC method (K-S test, *p*-value=0.345) and for the MR method (K-S test, *p*-value=0.586). This is the case neither for the DC nor for the SR methods (K-S test, *p*-values smaller than 10^*-*8^).

**Figure 4:**
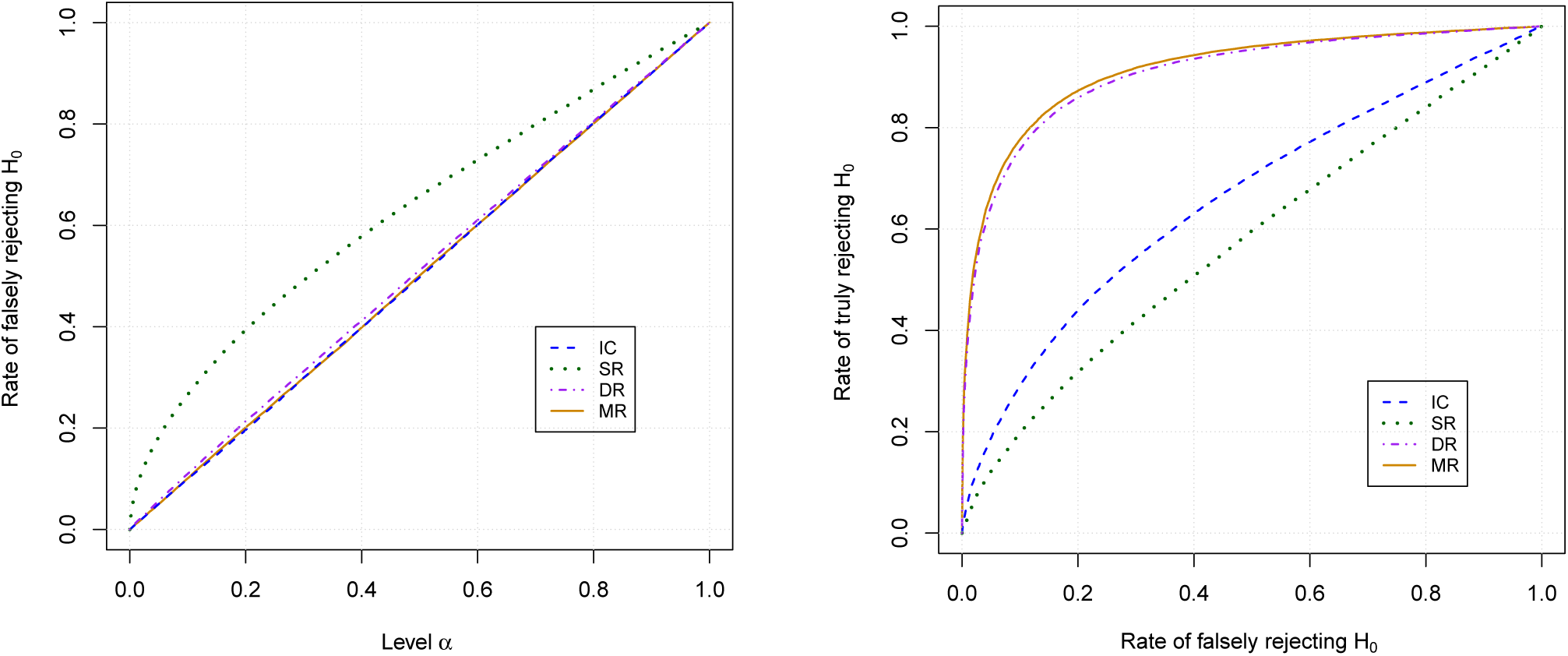
Left: Rates of false rejection of the null hypothesis at level *α vs α* when trait *A* follows the BM model with variance 0.09 and trait *B* follows the ABM model with trend 0.5 and variance 0.09. Right: ROC plots of the correlation tests obtained from two simulated traits under a BM model and an ABM model with trend 0.5 respectively, both with variance 0.09. Negative population is simulated with uncorrelated traits and positive population with traits correlated with correlation 0.5.

The ROC plots displayed in Figure 4-Right shows that performances of the IC and the SR tests are significantly lower than that of the MR and DC tests. The MR method is slightly more accurate than the DC test.

### 4.4 Correlation tests between two traits under directional evolution

In the case where the two traits are under directional evolution (here trait A has trend 0.5, and trait B has trend 1 both with variance 0.09), the rate of type I error of both the SR and the IC methods becomes maximal (Fig. 5-Left). In plain English, the SR and the IC method systematically reject the hypothesis that the traits are uncorrelated, even when they are uncorrelated. This behavior clearly prevents us to use the IC and the SR methods to detect correlation between traits under directional evolution. Still with regard to type I errors, performances of the MR and DC methods are essentially the same as in the case of neutral evolution or in the “mixed” case. The MR method looks perfect and the DC method still tends to reject the null hypothesis a little bit more than it should do. Taking a closer look on the *p*-values of the tests, we observe that those of the IC method no longer follow an uniform distribution (K-S test, *p*-value below 10^*-*9^) and so do those of the SR and DC methods. Only the MR method has the expected behavior of a test in this situation (K-S test, *p*-value=0.935).

**Figure 5:**
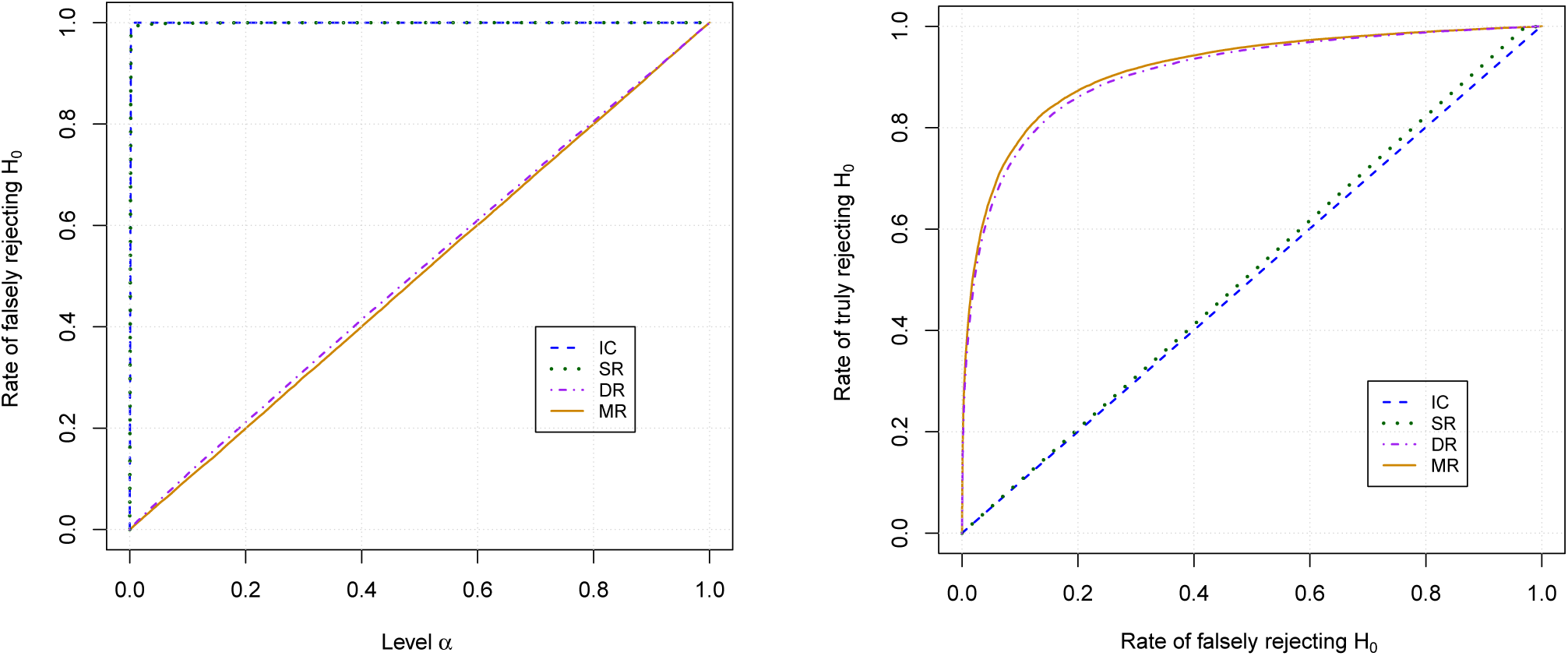
Left: Rates of false rejection of the null hypothesis at level *α vs α* when both traits *A* and *B* follow the ABM model with variance 0.09, and trend 0.5 and 1 respectively. Right: ROC plots of the correlation tests obtained from two simulated traits under ABM models with trend 0.5 and 1 respectively, both with variance 0.09. Negative population is simulated with uncorrelated traits and positive population with traits correlated with correlation 0.5.

Figure 5-Right displays the ROC plots of the SR, IC, DC and MR tests. The performance of the IC test is not better than a random guess and that of the SR test is almost as bad. The accuracy of the MR and DC tests is essentially the same as in the preceding case.

### 4.5 Discussion

As expected, the less accurate test is SR in all the situations. Overall, we observe that despite the flaws in its statistical properties, the DC method performs generally well with regards to the two criteria considered, whatever the trends of the traits. However, the DC method is always outperformed by the MR method. The “historical” IC method is outperformed by both the MR and the DC methods as soon as one of the traits is under directional evolution. It has the best performance from the ROC criteria only when the two traits evolve under the BM model, which is not very surprising since it corresponds exactly to the assumptions of this method, but the accuracy of the MR and DC methods is very close. The additional figures obtained from a greater variety of parameters and provided in the Supplementary Information lead to the same observations.

The simulations suggest to first test for the presence of a non-zero trend on each trait to compare, for instance by using the method of Appendix C, then to use the MR method if at least one of the traits shows a significant trend, and to use the standard IC method only if the two traits are under neutral evolution.

## 5 Correlation between hominin cranial capacity and body mass

### 5.1 Data

Evolution of hominin cranial capacity and body mass was studied in numerous works (Kappelman 1996; Henneberg 1998; Wood and Collard Leonard et al. 2003; Falk et al. 2005; Weber et al. 2005; Martin et al. 2006; Young 2006; Snodgrass et al. 2009; Montgomery et al. 2010; Potts 2011; Shultz et al. 2012; Schoenemann 2013; Hofman 2014; Grabowski et al. 2015; Grabowski 2016; Argue et al. 2017; Will et al. 2017; Du et al. 2018).

Our study is based on the hominin phylogenetic tree summarizing the best trees obtained in the dated Bayesian analysis of Dembo et al. (2015, Fig. 1), which is displayed in Figure 2. We combined data from several articles in order to get the body mass and the cranial capacity of as many species as possible, namely from Kappelman (1996, Table 1), Wood and Collard (1999, Table 3), Leonard et al. (2003, Table 3), Young (2006, Table 1), Schoenemann (2013, Tables 8.1 and 8.2), Grabowski et al. (2015, Table 4), Will et al. (2017, Table 4) and Du et al. (2018, Elec. Supp.). We excluded data associated to ambiguously identified species and to juvenile specimens. We finally averaged all the collected cranial capacities and body masses by species in order to obtain the data displayed in Table 1. We excluded *H. floresiensis* for calibrating our models, because being an outlier (Weber et al. 2005; Martin et al. 2006; Falk et al. 2007; Argue et al. 2017), this species over-influenced the results. Each time that a data required in an analysis was missing, we did not consider the corresponding species in this analysis. In particular, the correlation study pertains only to species for which both cranial capacity and body mass are known.

**Table 1:**
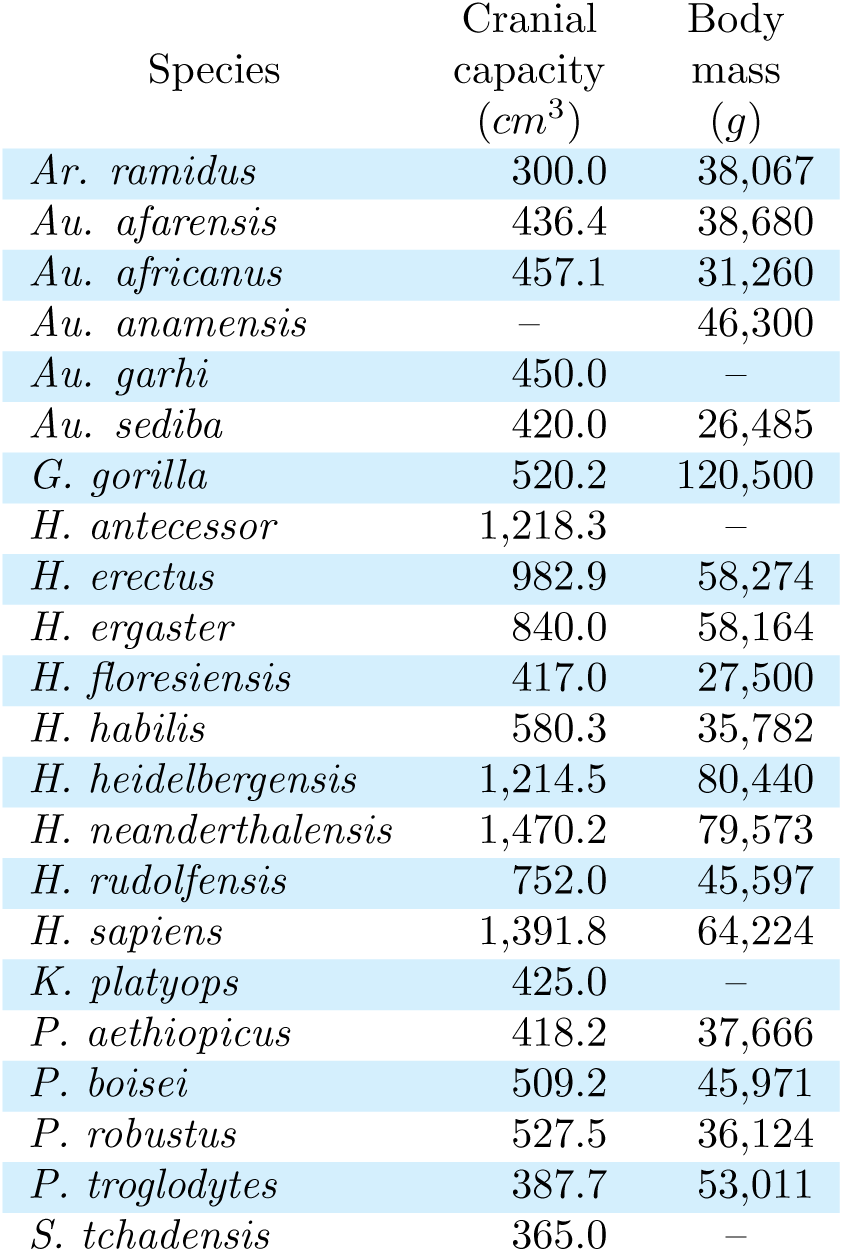
Cranial capacity and body mass of species of the phylogenetic tree of Figure 2.

| Species | Cranial<br>capacity<br>($cm^3$) | Body<br>mass<br>($g$) |
| --- | --- | --- |
| *Ar. ramidus* | 300.0 | 38,067 |
| *Au. afarensis* | 436.4 | 38,680 |
| *Au. africanus* | 457.1 | 31,260 |
| *Au. anamensis* | – | 46,300 |
| *Au. garhi* | 450.0 | – |
| *Au. sediba* | 420.0 | 26,485 |
| *G. gorilla* | 520.2 | 120,500 |
| *H. antecessor* | 1,218.3 | – |
| *H. erectus* | 982.9 | 58,274 |
| *H. ergaster* | 840.0 | 58,164 |
| *H. floresiensis* | 417.0 | 27,500 |
| *H. habilis* | 580.3 | 35,782 |
| *H. heidelbergensis* | 1,214.5 | 80,440 |
| *H. neanderthalensis* | 1,470.2 | 79,573 |
| *H. rudolfensis* | 752.0 | 45,597 |
| *H. sapiens* | 1,391.8 | 64,224 |
| *K. platyops* | 425.0 | – |
| *P. aethiopicus* | 418.2 | 37,666 |
| *P. boisei* | 509.2 | 45,971 |
| *P. robustus* | 527.5 | 36,124 |
| *P. troglodytes* | 387.7 | 53,011 |
| *S. tchadensis* | 365.0 | – |

We considered the logarithms of cranial capacity and body mass data such as in Kappelman (1996); Henneberg (1998); Leonard et al. (2003); Snodgrass et al. (2009); Navarrete et al. (2011); Du et al. (2018). Taking the logarithm of quantitative trait values is quite usual since it accounts for the fact that for instance, an increase of 100 g does not have the same significance for an organism of 1 kg as for a organism of 100 kg. From a statistical point of view, log-transformation is a particular case of Box-Cox transformations which tend to stabilize the variance. It is also sometimes used to approach Gaussian behavior required by Brownian evolution models (Legendre and Desdevises 2009).

We applied diagnostic tests on residuals after log-transformation in order to check for least squares regression validity conditions. Namely, we use Durbin-Watson’s test to detect autocorrelation at lag 1 (Durbin and Watson 1950, 1951, 1971); Harrison-McCabe’s test to detect heteroscedasticity (Harrison and McCabe 1979) and Jarque-Bera’s test to confirm normality (Jarque and Bera 1987).

### 5.2 Evolutionary trends of hominin cranial capacity and body mass

Several works agree with the fact that hominin brain size increased through evolution (Henneberg 1998; Montgomery et al. 2010; Navarrete et al. 2011; Potts 2011; Shultz et al. 2012; Hofman 2014). Henneberg (1998) found a significant correlation between the log-transformed cranial capacity and the fossil age through a direct “non-phylogenetic” regression approach. Applying the same approach on our dataset, we also detected a positive evolutionary trend (*p*-value=0.003), but least squares regression conditions are violated (Harrison-McCabe’s test, *p*-value=0.001), certainly because of the phylogenetic relationships between tips values. Conversely, the hominin cranial capacities fulfill the conditions of the phylogenetic trend detection test presented introduced in Appendix C (Durbin-Watson’s test, *p*-value=0.422; Harrison-McCabe’s test, *p*-value=0.604; Jarque-Bera’s test, *p*-value=0.646). The test of Appendix C concludes to a positive trend (*p*-value=0.012) which amounts to multiplying the cranial capacity by 1.2 per Ma.

Several studies conclude that the body mass data increased during evolution by using non phylogenetic approaches, i.e., without taking into account the evolutionary relationships between the species (Henneberg 1998; Will et al. 2017). Considering a direct “non phylogenetic” regression of the logarithm of the body mass of our dataset led to detect a positive evolutionary trend (*p*-value=0.002), whereas, by taking into account the evolutionary relationships between species, the detection test of Appendix C did not conclude to a significant trend on the logarithm of hominin body mass (*p*-value=0.072). Both the non phylogenetic regression and our detection test satisfy the regression assumptions (Durbin-Watson’s test, *p*-value=0.243 and *p*-value=0.227; Harrison-McCabe’s test, *p*-value=0.066 and *p*-value=0.796; Jarque-Bera’s test, *p*-value=0.953 and *p*-value=0.865 respectively).

Our results are consistent with those of Montgomery et al. (2010), who also found a positive trend in cranial capacity and no significative trend in the body size evolution of hominids. Finally, testing for correlation between the logarithms of hominin cranial capacity and body mass falls in a situation close to that of Section 4.3, in which we compared a trait simulated under neutral evolution with a trait simulated under directional evolution.

### 5.3 Correlation between hominin cranial capacity and body mass

We applied the MR method in order to test for correlation between the logarithms of the hominin body mass and cranial capacity. These data fulfill the requirements of our correlation test (Durbin-Watson’s test, *p*-value=0.819; Harrison-McCabe’s test, *p*-value=0.690; Jarque-Bera’s test, *p*-value=0.461). Durbin-Watson’s and Harrison-McCabe’s tests require to order the residuals with respect to the explanatory variable, which is straightforward for single regressions, but not for multiple regressions. Following Fan and Huang (2001), we ordered the residuals with respect to most informative linear combination of the explanatory variables obtained from principal component analysis.

It is thus allowed to apply the MR test, which concluded that the logarithms of hominin cranial capacity and body size are significantly correlated (*p*-value=0.027). In plain English, even by taking into account possible evolutionary trends, the logarithms of these two traits do not change independently. This result is consistent with Grabowski (2016), who showed that the evolution of hominin body size and that of the cranial capacity are related.

## A Distribution of independent contrasts – Proof of Theorem 3

Let us first prove by induction that for all nodes *k* of *𝒯*, we have that

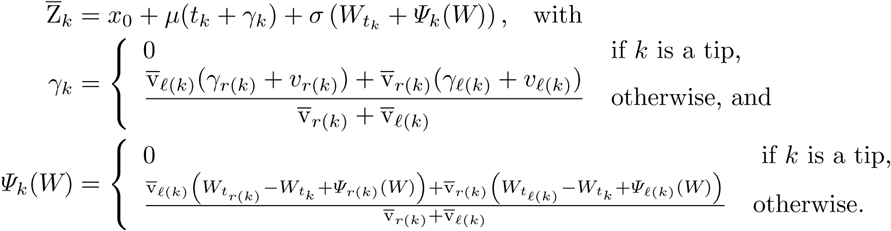

In the base case, i.e., when *k* is a tip, the property is granted since from Definition 2, we have that

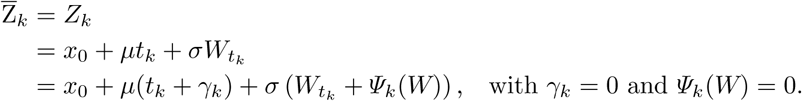

Let *k* be an internal node and let us assume that the property holds for its direct descendants *r*(*k*) and *ℓ* (*k*). From Definition 2, we have that

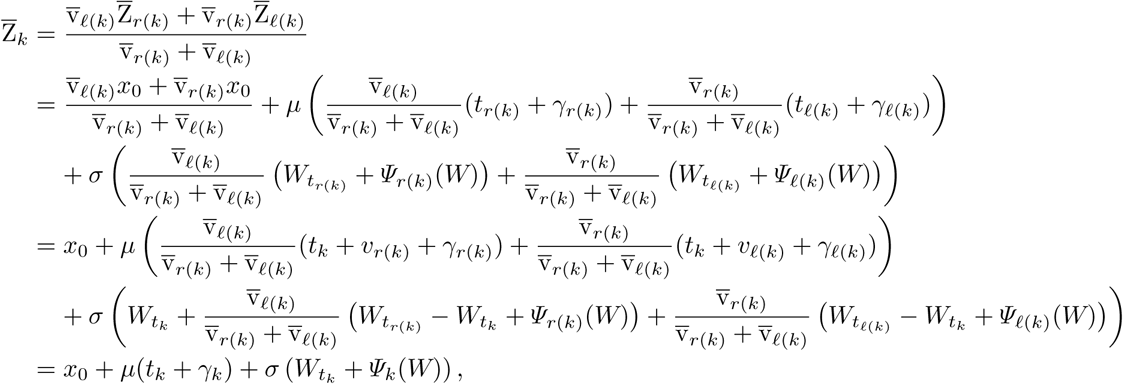

by setting

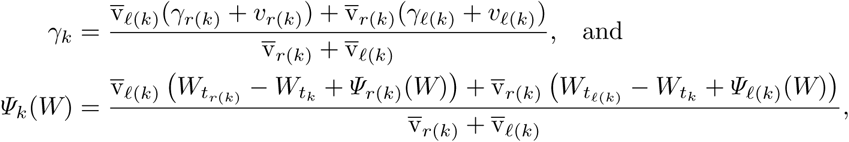

and the property holds for all nodes *k* of 𝒯.

Proving the second point of the theorem is direct since from Definition 2, we have that

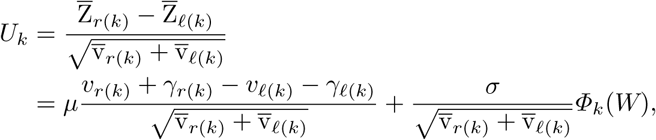

where

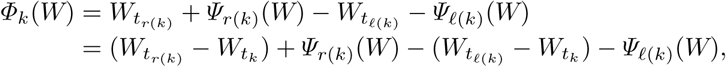

By construction, for all nodes *k, Ψ*_*k*_(*W*) is a linear combination of independent centered Gaussian random variables of the form 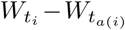 where *i* is a descendant of *k*. It follows that *Ψ*_*k*_(*W*) is a centered Gaussian variable, which is independent from any random variable 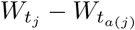 if *j* is not a descendant of *k*. Since (*W*_*t*_)_*t*>0_ is the Wiener process, we have that

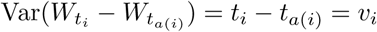

for all nodes *i* of *𝒯* (Grimmett and Stirzaker 2001).

Let us prove by induction that

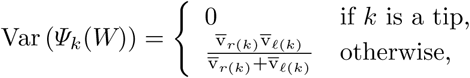

which is equivalent to say that Var 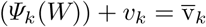 from Definition 2. It is basically true in the base case where *k* is a tip. If *k* is an internal node, by assuming that the induction assumption holds for its direct descendants *r*(*k*) and *ℓ* (*k*), we have that

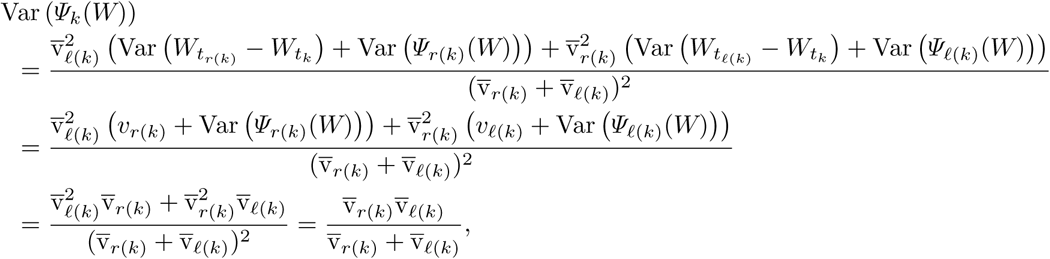

which proves the form of Var (*Ψ*_*k*_(*W*)).

In the same way, *Φ*_*k*_(*W*) is a linear combination of independent centered Gaussian random variables of the form *W*_*i*_ − *W*_*a*(*i*)_ where *i* is a descendant of *k*, with variance

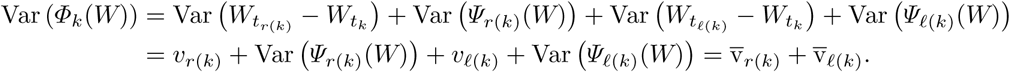

## B Ultrametric trees - Proof of Proposition 5

Let *𝒯* be an ultrametric tree and *T* be the total path-length/time from its root to its tips.

We shall prove by induction that *γ*_*k*_ = *T* −*t*_*k*_ and *h*_*k*_ = 0 for all nodes *k* of. 𝒯 The property is basically true if *k* is a tip, our base case. Let *k* be an internal node and let us assume that the property holds for its direct descendants *r*(*k*) and *ℓ*(*k*). From Theorem 3, we have that

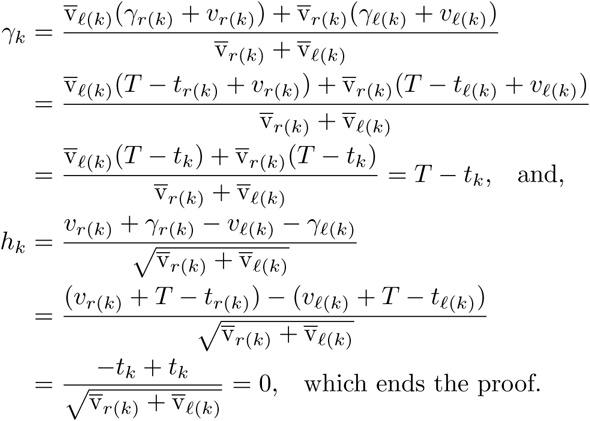

## C Trend estimation and detection

Equation 1 shows that under an ABM model with trend *µ*, the independent contrasts *U*_*k*_ can be written as the product of *µ* with the corresponding temporal variable *h*_*k*_, plus an independent, centered Gaussian term of constant variance with respect to *k*. This suggests to estimate the trend *µ* as the slope of the following linear equation

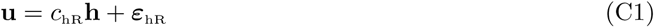

where, from Equation 1, the entries of the error vector *ε*_hR_ are samples of random variables

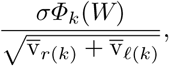

which are independent and Gaussian distributed with mean zero and variance *σ*^2^ under the current assumptions.

From Equation C1, the linear regression estimator of *µ* is

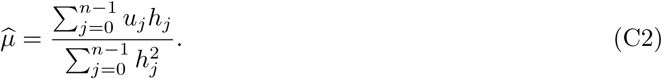

Again from Equation C1, testing for a trend in the evolution of trait under an ABM model can be performed by testing for the nullity of the slope. In order to assess the accuracy of this trend test (referred to as the “hR test”) with regard to the standard regression of the tips values with respect to their times (referred to as the “SR test”), we simulated evolution of quantitative traits with and without trend on the tree of Figure 2 and plot the type I error rate against the rejection level and the ROC plots of these two tests. Results are displayed in Figure C1. The hR test clearly outperforms the SR test. Figure C1-Left shows that the SR test rejects the null hypothesis more than it should do whereas the hR test rejects it at the exact level required. Moreover, their ROC plots show that the hR test better discriminates between traits simulated with and without trend than the SR test (Fig. C1-Right).

**Figure C1:**
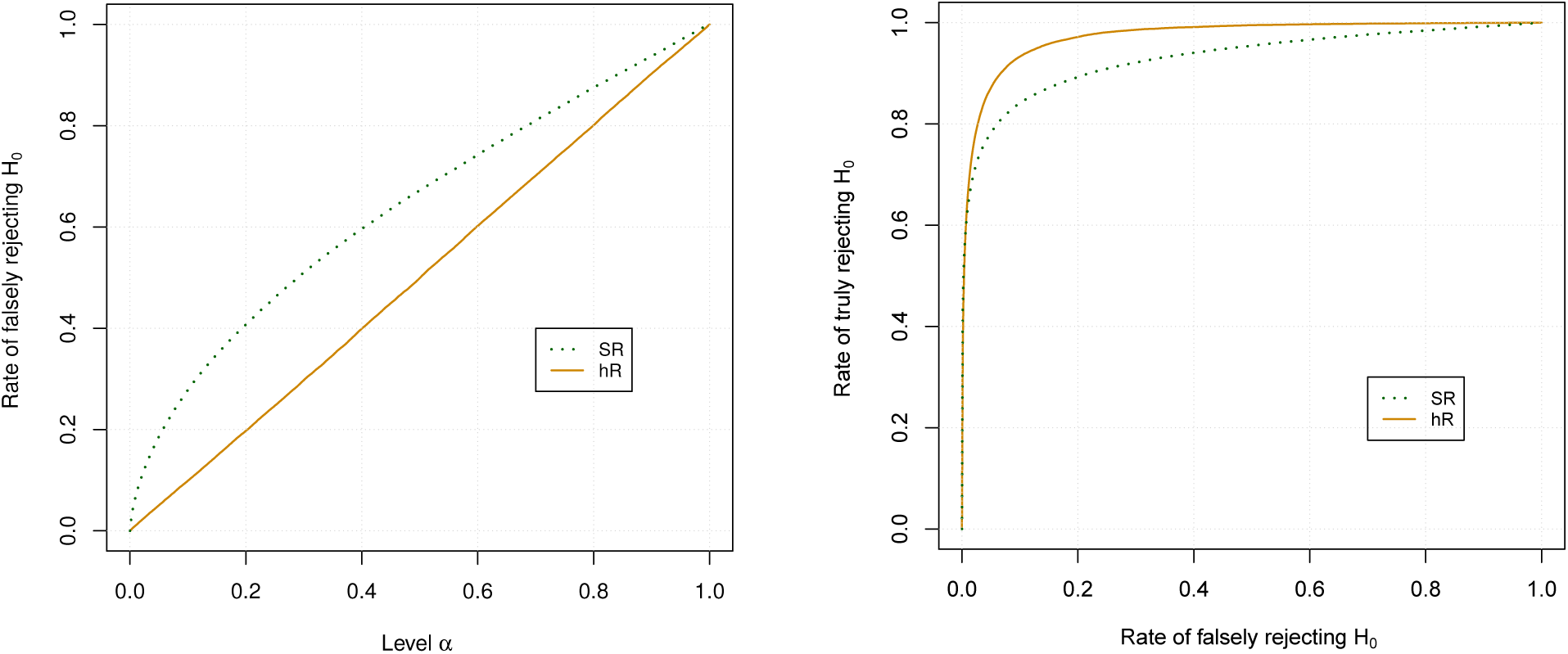
Left: Rates of false rejection of the null hypothesis (i.e., no trend) at level *α vs α* on traits simulated under the BM model with variance 0.09. Right: ROC plots of the trend detection tests obtained from simulated traits under the BM model for negative population (i.e., under H_0_) and under the ABM model with trend 0.2 for the positive population, both with variance 0.09.

## D Directional contrasts

### Equivalence between centered and directional contrasts

Elliott (2015) associated to all nodes *k* of *𝒯* and all values *β* the quantity *e*_*k*_(*β*) defined as

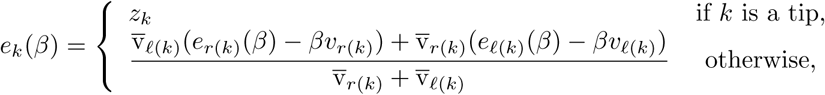

where the modified branch lengths 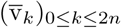 are given in Definition 1.

For all internal nodes *k* of *𝒯*, Elliott (2015) then defined the *(β-)directional contrast* as

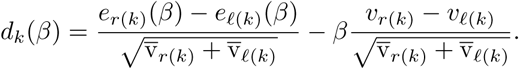

Let us start to prove by induction that for all nodes *k* of *𝒯*, we have that 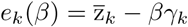.

The equality is basically true in the base cases, since if *k* is a tip of *𝒯*, we have that *γ*_*k*_ = 0 and 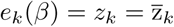.

Let *k* be an internal node of 𝒯 and let us assume that the equality holds for its two direct descendants *r*(*k*) and *ℓ*(*k*). We have

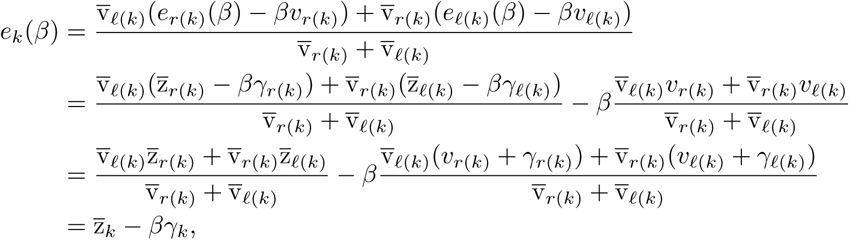

which proves the equality for all nodes of *𝒯*.

Last, we have that

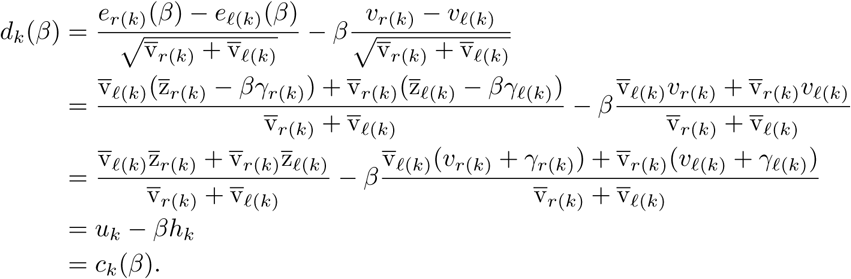

In plain English, for all internal nodes *k* and all values *β*, the *β*-directional contrast *d*_*k*_(*β*) is equal to the *β*-centered contrast *c*_*k*_(*β*).

### Estimated directional contrasts

In order to correct the trend effect, Elliott (2015) proposed to consider the 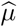-directional contrasts where 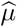 is the estimated trend (Equation C2).

Let *B* be the random variable associated to the estimated trend. From Equation C2, we have that

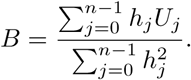

From Corollary 4, if the trait follows the ABM model of parameters (*x*_0_, *µ, σ*^2^), then for all internal nodes *k* of *𝒯*, the independent contrast random variables *U*_*k*_ are Gaussian distributed with

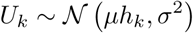

which implies that *B* follows the Gaussian distribution 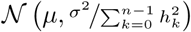 under the ABM model (*x*_0_, *µ, σ*^2^).

For all internal nodes *k* of 𝒯, the random variable *D*_*k*_(*B*) associated to the *k*^th^ estimated directional contrast is defined as

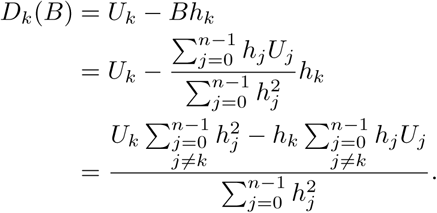

Corollary 4 implies that the estimated directional contrast random variables *D*_*k*_(*B*) are Gaussian distributed with

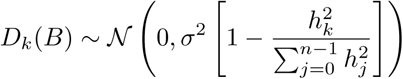

Moreover, since the independent contrast random variables *U*_*k*_ are independent from one another, we have that for all *i* ≠ *k*

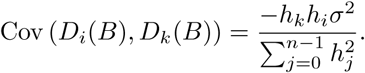

Since *h*_*k*_ ≠ 0 in the general case, the estimated directional contrast random variables are neither identically distributed nor independent.

## E IC and PGLS regressions – Proof of Theorem 8

In all what follows, **0** (resp. **1**) denotes the column vector with all entries equal to 0 (resp. to 1; their dimensions depending on the context) and for all numbers *N*, **I**_*N*_ is the identity matrix of dimension *N* × *N*. The transpose of a matrix or a vector *A* is noted *A*^*′*^. We recall that *n* is the number of internal nodes of *𝒯* which thus has *n* + 1 tips.

### A matrix presentation of the Felsenstein’s (1973) algorithm

Let us sketch a matrix presentation of the Felsenstein’s (1973) algorithm which iteratively computes the following variables for all nodes *k* of 𝒯. By putting 𝒯_*k*_ for the subtree of 𝒯 rooted at *k* and *n*_*k*_ for its number of internal nodes, let

- **g**_*k*_ be the vector of dimension *n*_*k*_ + 1 that is such that the “artificial” trait value of node *k*, i.e., 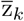 of Definition 1, is obtained by multiplying 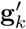 with the tip value vector of *𝒯*_*k*_,
- *Q*_*k*_ be the *n*_*k*_ × (*n*_*k*_ + 1) matrix giving the independent contrasts of the subtree 𝒯_*k*_ from its tip value vector, and,
- *δ*_*k*_ be the increment applied to the branch ending by *k* (i.e., 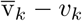 in Definition 1).

By initializing g_*k*_ to the vector [1], *Q*_*k*_ to the 0 ×0 “empty” matrix and *δ*_*k*_ to 0 for all tips *k* of 𝒯, these variables are recursively computed for all internal nodes *k* of 𝒯 with direct descendants *ℓ* (*k*) and *r*(*k*) by setting:

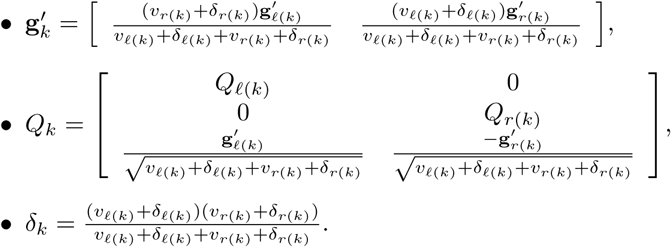

It is straightforward to prove by induction that 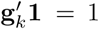 for all nodes *k* of *𝒯*. Let us prove by induction that 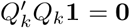 for all nodes *k* of 𝒯 The property is basically true for all tips. Let us assume that *k* is an internal node and that its direct descendants *ℓ*(*k*) and *r*(*k*) both satisfy the property. We have

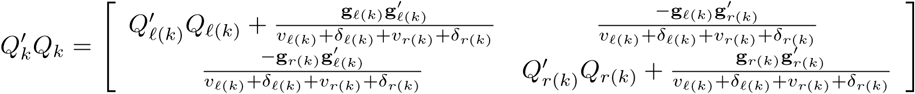

thus

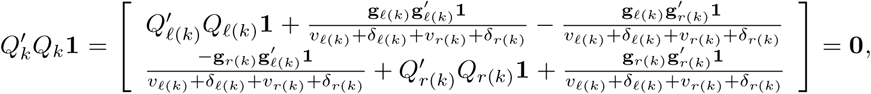

since 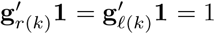 from above and 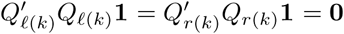 from the induction hypothesis, which proves that 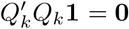 for all nodes *k* of 𝒯

Let us set 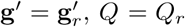 and *δ* = *δ*_*r*_ where *r* is the root of 𝒯 and let ∑ be the covariance matrix associated to 𝒯, i.e., for all pairs of tips (*i, j*) of 𝒯, the (*i, j*)-entry of ∑ is the total time between the root and the most recent common ancestor of *i* and *j*. Felsenstein (1973) showed that by assuming that a trait follows a Brownian process with variance *σ*^2^ and by putting **Z** for the random vector of its tip values, the contrasts, i.e., the entries of *Q***Z**, are independent centered Gaussian variables with variance *σ*^2^ and that **g**^*′*^**Z** is a centered Gaussian variable with variance *σ*^2^*δ* which is independent from all the contrasts. It follows that if **Z** is a Gaussian vector with covariance matrix *σ*^2^∑ then *Q***Z** is a *n*-Gaussian vector with covariance matrix *σ*^2^**I**_*n*_ and, by defining the (*n* + 1) ×(*n* + 1) matrix 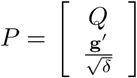, that *P***Z** is a (*n* + 1)-Gaussian vector with covariance matrix *σ*^2^**I**_*n*+1_. Since *P* is invertible (the standard presentation of the Felsenstein’s (1973) algorithm shows that *P* can be written as a product of invertible matrices), this implies *P*′*P* = ∑^*-*1^ and that

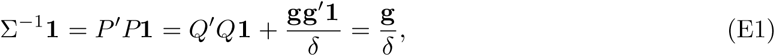

which itself directly implies that

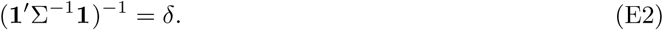

Moreover, since ∑^*-*1^ is symmetric, Equation E1 implies that **gg**′ = *δ*^2^∑^*-*1^**11**^*′*^∑^*-*1^ and that

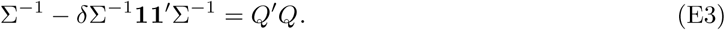

Note that matrix *P* plays the same role as the *phylogenetic transformation matrix* in Adams and Collyer (2015) and as the matrix **D** defined in a complete different way in Garland and Ives (2000, p349).

### Multiple regression with IC and PGLS

Let **y** and *X* be respectively a vector of tip values of a trait and a matrix of tip values of *p* traits or co-variables (e.g., the time, an environmental variable etc.). We shall show that testing for correlation between **y** and any column of *X* leads to the same result with the IC and the PGLS methods.

In the IC regression and under the notations of subsection above, we consider the following linear equation

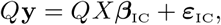

where the error vector ***ε***_IC_ is a realization of the centered Gaussian random vector with covariance matrix proportional to **I**_*n*_. Vector *Q***y** has dimension *n* and there are *p* regressors. The vector of estimated regression coefficients is

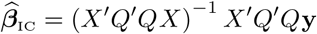

and the estimator variance of the *i*th coefficient 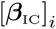 is 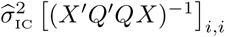 where the residual variance estimate 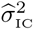 is

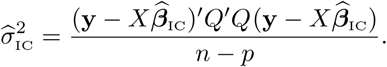

Under the key regression assumptions and that the *i*th coefficient 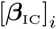 is null, the statistics

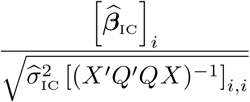

follows the Student distribution with *n* − *p* degrees of freedom.

In the PGLS regression, the linear equation is

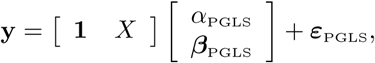

where the error vector is a realization of the centered Gaussian vector with covariance matrix proportional to ∑ = (*P* ′*P*)^*-*1^. Vector **y** has dimension (*n* + 1) and there are *p* + 1 regressors (including the intercept). The vector of estimated regression coefficients is

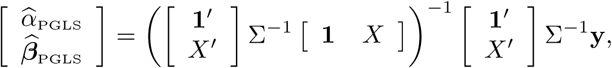

and the estimator variance of the *i*th coefficient 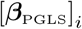 is

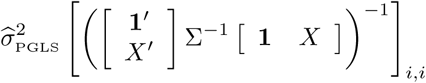

where the residual variance estimate 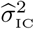 is

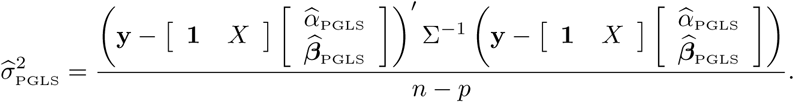

Under the regression assumptions and that the *i*th coefficient 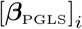 is null, the statistics

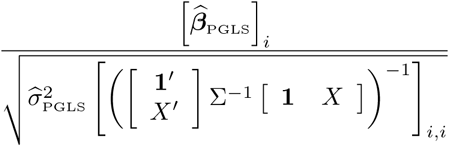

follows the Student distribution with *n* − *p* degrees of freedom.

In order to prove that the IC and the PGLS methods are equivalent to test for correlation in a multiple regression context, we shall establish that the three following properties hold:

1. 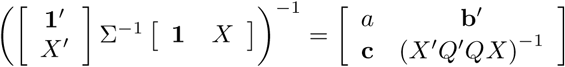 for a real *a* and two *n*-vectors **b** and **c**,
2. 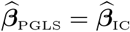,
3. 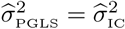.

From the block matrix inversion formula and Equations E1, E2 and E3, we get that

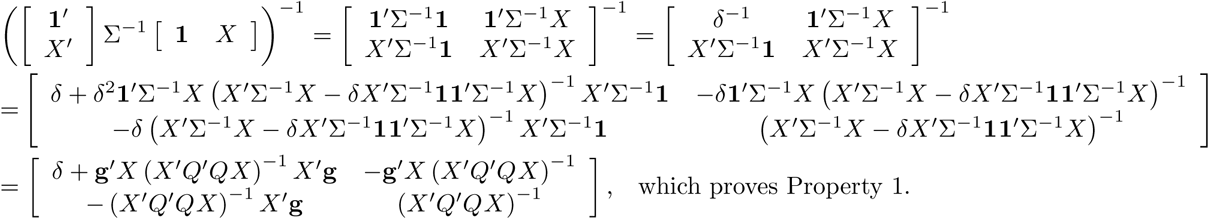

The vector 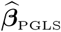 of the PGLS regression coefficient estimates without the intercept is obtained by multiplying the second line of the block matrix above with the column 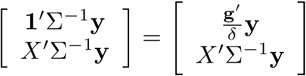:

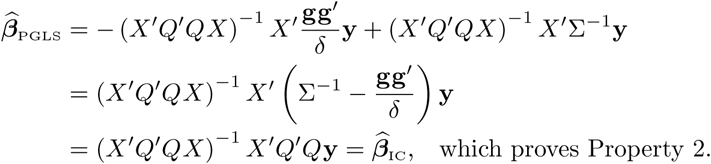

In the same way, the intercept estimate of PGLS is

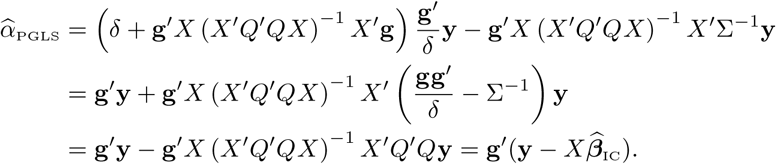

The residual variance estimate of PGLS is given by

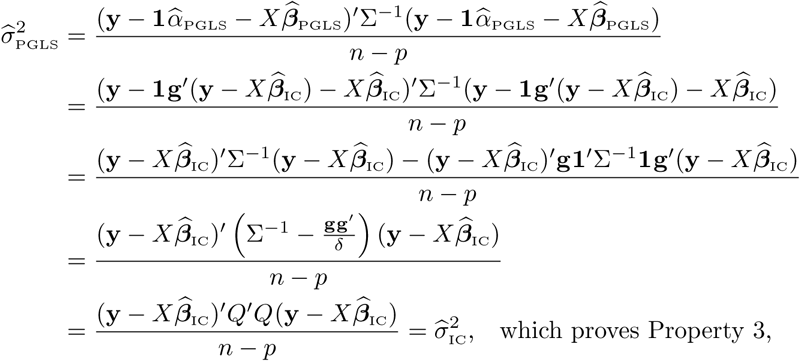

and ends to prove that testing for correlation with IC and PGLS is equivalent.

### Multiple regression with time as co-variable

Let us start by showing that for all internal nodes *k, h*_*k*_ is the phylogenetic contrast of the tip times associated to *k*. By putting 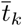 for the “artificial” time reconstructed at the internal node *k* by the Felsenstein’s (1973) algorithm, it is straightforward to prove by induction that 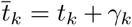 for all nodes *k* of *𝒯*. For all internal nodes *k* with direct descendants *r*(*k*) and *ℓ*(*k*), we have that

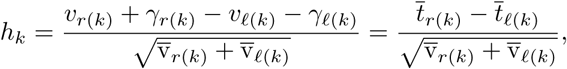

which is well the contrast of the tip times associated to node *k*. It follows that the vector **h** = (*h*_*k*_)_1≤*k<n*_ is obtained by multiplying the vector **t** = (*t*_*k*_)_*n*≤*k*≤2*n*_ of tip times by *Q*, i.e., **h** = *Q***t**.

Let us consider two traits *A* and *B* and their tip-value vectors 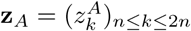 and 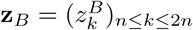.

The linear equation

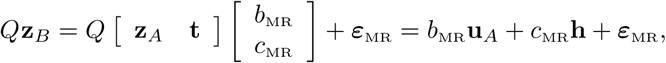

where the error vector *ε*_MR_ is a realization of a centered Gaussian vector with covariance matrix proportional to **I**_*n*_, corresponds to the regression considered in the MR method (Equation 8). We showed in the section above that testing for correlation between traits *A* and *B* with this equation is equivalent to testing for correlation between *A* and *B* with the linear equation

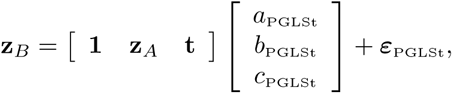

where the error vector *ε*_PGLSt_ is a realization of a centered Gaussian vector with covariance matrix proportional to ∑ which corresponds to testing for correlation between *A* and *B* with tip times as covariables by the PGLS method.

