## Additional figures for "Testing for correlation between traits under directional evolution" (from another version of this preprint: version 1, 2019-03-04)

### SI-1 Additional figures

We provide here additional plots of Type I error (Fig. SI-1) and ROC curves obtained from simulating two traits either uncorrelated or correlated with correlation 0.5 (Fig. SI-2) or correlated with correlation 0.7 (Fig. SI-3), under BM and ABM models with various trends.

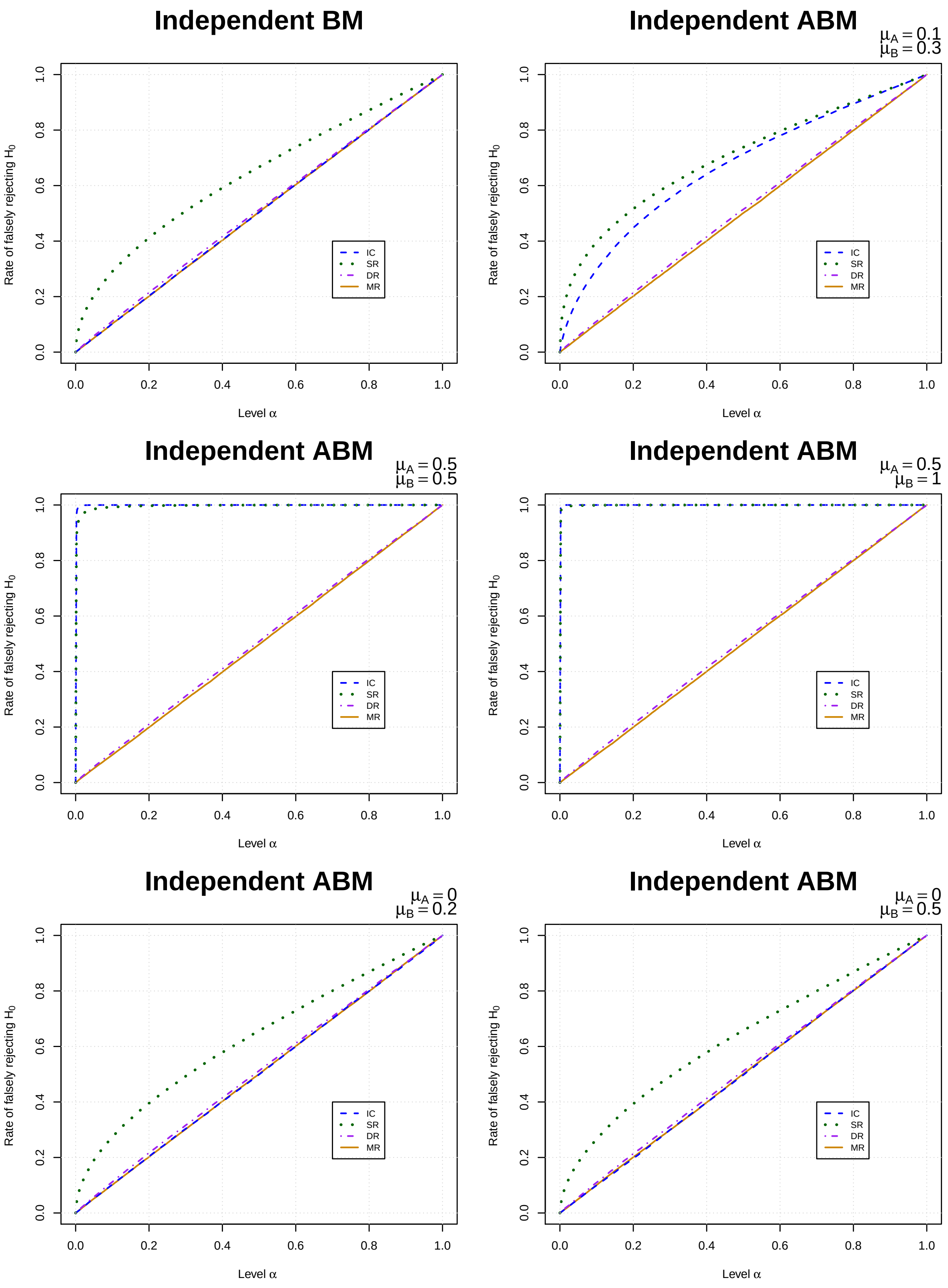


Figure SI-1: Rates of false rejection of the null hypothesis at level $\alpha$ *vs* $\alpha$ from two uncorrelated traits $A$ and $B$ simulated under various trends with variance 0.09.

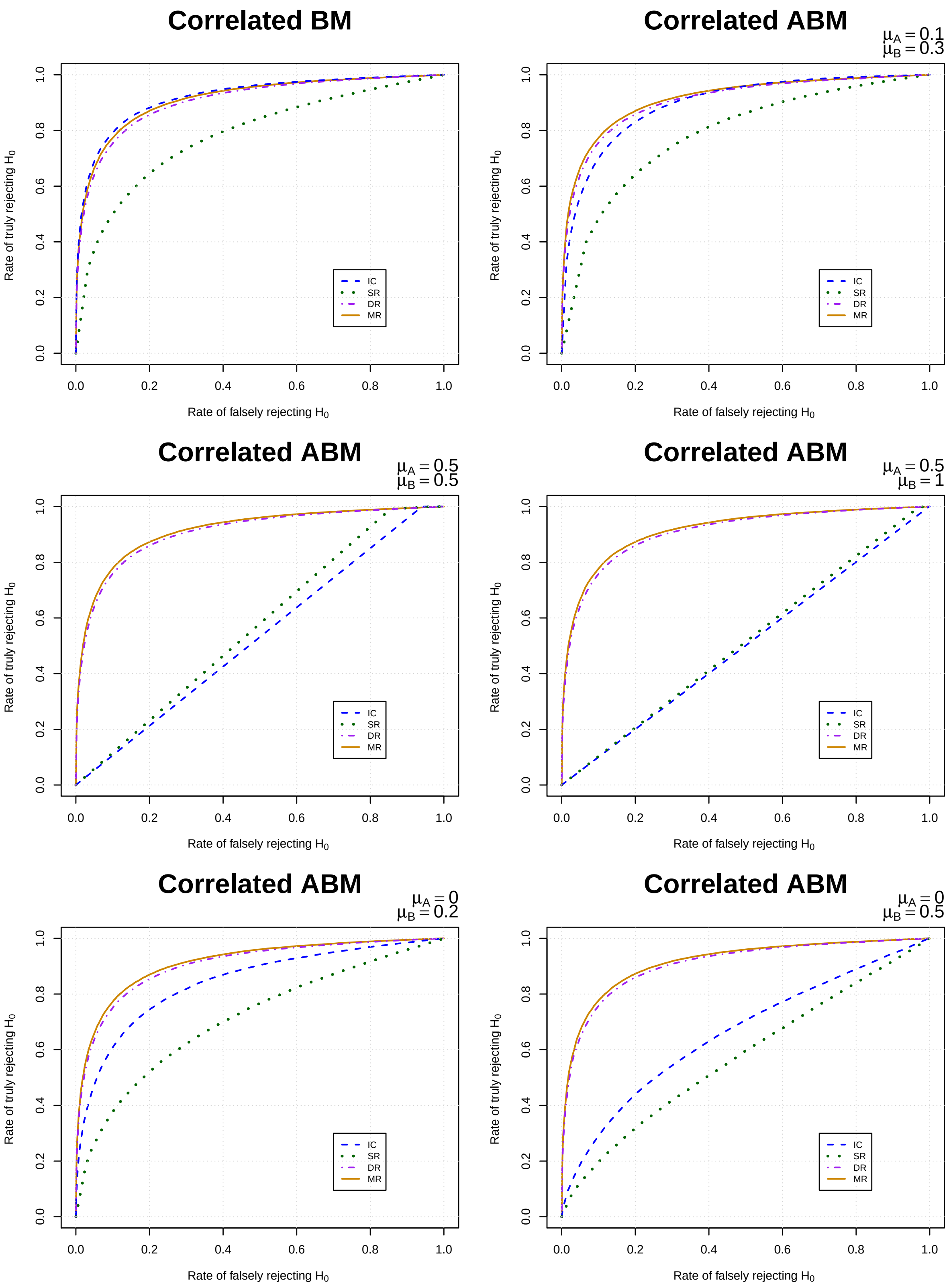


Figure SI-2: ROC plots of the correlation tests obtained from two traits $A$ and $B$ uncorrelated and correlated with correlation $\rho = 0.5$, simulated under various trends with variance 0.09.

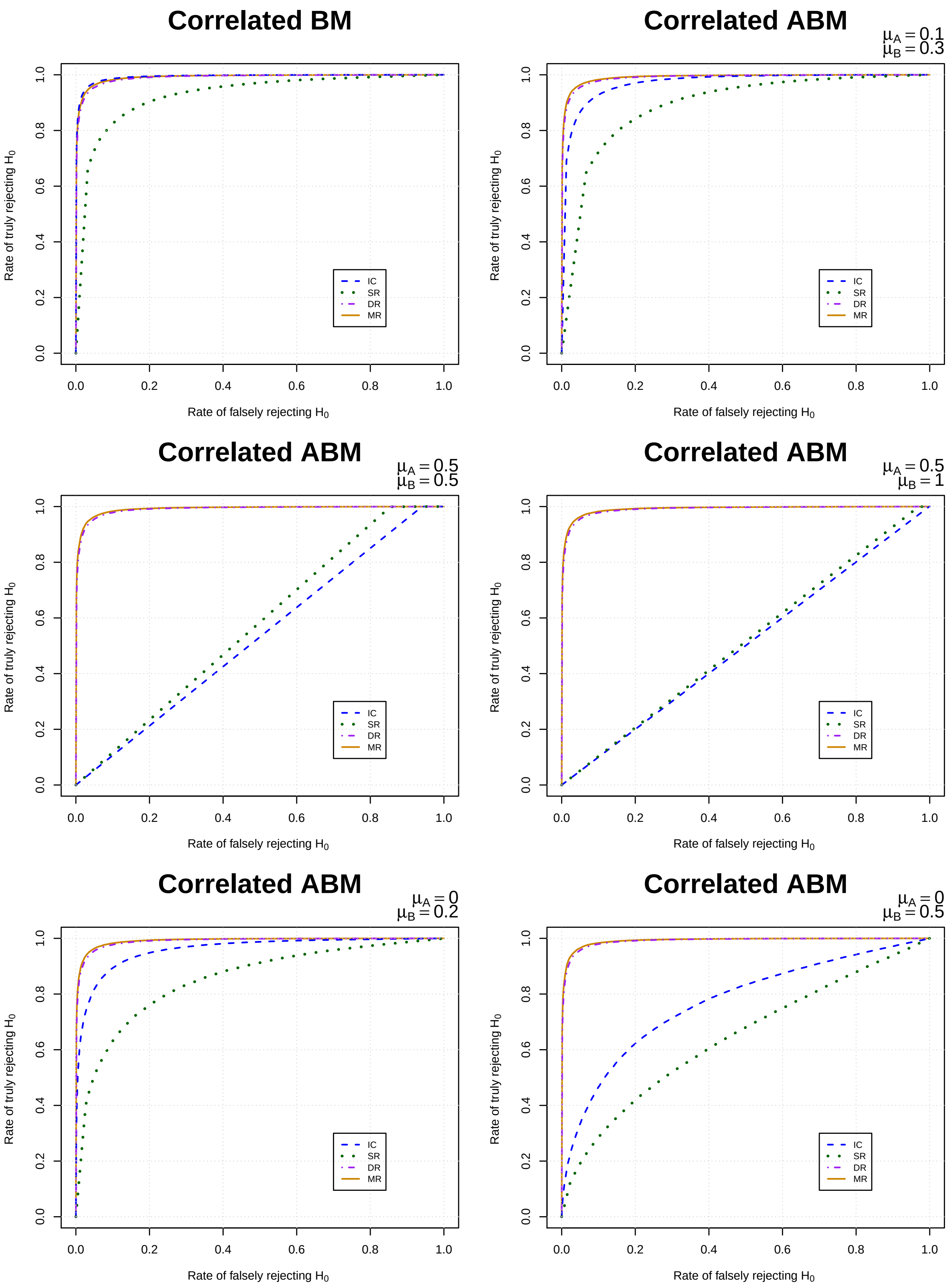


Figure SI-3: ROC plots of the correlation tests obtained from two traits $A$ and $B$ uncorrelated and correlated with correlation $\rho = 0.7$, simulated under various trends with variance 0.09.
